# Meiotic DNA repair in the nucleolus employs a non-homologous end joining mechanism

**DOI:** 10.1101/553529

**Authors:** Jason Sims, Gregory P. Copenhaver, Peter Schlögelhofer

## Abstract

Ribosomal RNA genes are arranged in large arrays with hundreds of rDNA units in tandem. These highly repetitive DNA elements pose a risk to genome stability since they can undergo non-allelic exchanges. During meiosis DNA double strand breaks (DSBs) are induced as part of the regular program to generate gametes. Meiotic DSBs initiate homologous recombination (HR) which subsequently ensures genetic exchange and chromosome disjunction.

In *Arabidopsis thaliana* we demonstrate that all 45S rDNA arrays become transcriptionally active and are recruited into the nucleolus early in meiosis. This shields the rDNA from acquiring canonical meiotic chromatin modifications, meiotic cohesin and meiosis-specific DSBs. DNA breaks within the rDNA arrays are repaired in a RAD51-independent, but LIG4-dependent manner, establishing that it is non-homologous end joining (NHEJ) that maintains rDNA integrity during meiosis. Utilizing ectopically integrated rDNA repeats we validate our findings and demonstrate that the rDNA constitutes a HR-refractory genome environment.

## Introduction

All organisms depend on well-maintained genetic information and its coordinated interpretation during their life cycle. DNA damage, if not reliably repaired, may lead to loss of genetic information, genomic rearrangements or cell cycle arrest. The most deleterious DNA insults are DNA double strand breaks (DSBs), which may occur through exposure to genotoxic agents, or as a result of errors during endogenous processes like DNA replication or meiotic recombination (Ceccaldi et al. 2016). DSBs can either be repaired by non-homologous end joining (NHEJ) or homologous recombination (HR) DNA repair pathways (Ceccaldi et al. 2016). Regardless of what pathways are used, repair of repetitive DNA, including the genes encoding ribosomal RNAs (rDNA), must be carefully regulated in order to prevent ectopic interactions.

NHEJ is believed to work during all cell cycle stages while HR appears to be the most prominent DNA repair pathway during S and G2 phases (Ceccaldi et al. 2016).

NHEJ pathways are further differentiated depending on the proteins involved. Canonical NHEJ (c-NHEJ) initiates with the recognition of DSBs by the conserved KU70/KU80 heterodimer that recruits, either directly or indirectly, its accessory factors including DNA-PKs, X-ray cross complementing protein 4 (XRCC4), XRCC4-like factor (XLF), and Apartaxin- and-PNK-like factor (APLF). These factors are required to stabilize and process the break to produce ligation competent DNA ends. Ligation of the processed ends is mediated by DNA Ligase IV, which is stabilized at the DSB by XRCC4 (Bennardo et al. 2008; Chiruvella et al. 2013; Davis and Chen 2013).

At least one alternative NHEJ pathway, called microhomology-mediated end joining (MMEJ) has been described and is characterized by requiring resected ends in order to function properly (McVey and Lee 2008). The resection of the ends is mediated by the MRE11-RAD50-NBS1/XRS2 (MRN/X) complex together with SAE2/COM1/CtIP and relies on PARP1, PARP2 and Ligase 3/XRCC1 for re-annealing the broken DNA ends (Dueva and Iliakis 2013). All of the afore mentioned processes and proteins are present in plants with the exception of DNA-PKcs, a Phosphatidyl inositol 3-kinase related kinase family member, (Templeton and Moorhead 2005; Yoshiyama et al. 2013; Shen et al. 2017) indicating that NHEJ may be differently regulated in plants.

The common denominator of all NHEJ pathways is direct ligation of the processed DNA ends, with the deleterious potential of losing or adding genetic information at the junction site or of connecting different chromosome arms, leading to translocations or losses (McVey and Lee 2008; Chang et al. 2017).

HR, in contrast, utilizes a repair template, either from the sister chromatid or the corresponding homologue, copying missing information that may have been lost during DNA damage or processing (Ceccaldi et al. 2016). HR is described as an error-free DNA repair pathway, yet it may lead to sequence deletions, duplications, inversions or translocations if the DNA lesion is misaligned with a non-allelic template during repair (Sasaki et al. 2010).

A special case of HR takes place during meiosis. Meiosis is a pair of specialized cell divisions used by sexually reproducing organisms to produce generative cells. During meiosis the genome is reduced by half and paternal and maternal genetic information is recombined to yield novel allelic combinations along each chromosome. Meiotic recombination is initiated by formation of enzymatically mediated DSBs. These are catalysed by the conserved topoisomerase-like complex SPO11/MTOPVIB (Keeney et al. 1997; Hartung et al. 2007; Borde and de Massy 2013; Vrielynck et al. 2016), in conjunction with other partners (Robert et al. 2016), and are a prerequisite for subsequent crossover formation (Hunter 2015; Mercier et al. 2015). During the cleavage reaction, two SPO11 moieties become covalently linked to the two 5′ ends of the DSB. DSBs are further processed by the conserved MRN/X complex together with COM1/SAE2/CtIP, which release the SPO11 proteins attached to a short DNA oligonucleotide (Keeney and Kleckner 1995; Prinz et al. 1997; Neale et al. 2005; Prieler et al. 2005; Uanschou et al. 2007). Further resection of the 5′ end by 5′-3′ exonucleases extends the single-stranded 3′ tails (Garcia et al. 2011; Symington 2014). These overhangs are bound by ssDNA binding proteins and subsequently loaded with RAD51 and meiosis specific DMC1 recombinases (Shinohara et al. 1997; Kurzbauer et al. 2012; Da Ines et al. 2013; Brown and Bishop 2014). Meiotic DSBs are repaired by HR, but a dedicated set of proteins, among them DMC1 and ASY1/HOP1/HORMAD1/2 (Caryl et al. 2000; Niu et al. 2005; Wojtasz et al. 2009), biases DNA repair to use homologous sequences on non-sister chromatids of the homologous chromosome (Caryl et al. 2000; Petukhova et al. 2003; Kerzendorfer et al. 2006; Sanchez-Moran et al. 2007; Kim et al. 2010; Uanschou et al. 2013; Stronghill et al. 2014). Of the 170 - 350 DSBs introduced per meiocyte (numbers range from yeast to mouse; *A. thaliana* Col-0 has about 250 breaks per cell (Kurzbauer et al. 2012)), only a few mature into crossovers (about 10 per meiocyte in *A. thaliana* (Armstrong and Jones 2003)). The remaining DSBs are repaired as non-crossovers via different sub-branches of the HR pathway (reviewed in Mercier et al. 2015). A key feature of all HR DNA repair processes is the precise alignment of DSB sites with their corresponding repair templates which occurs via the formation of the axis and synaptonemal complex (SC) that include proteins such as ASY1, ASY3, ASY4 and ZYP1 (Storlazzi et al. 1996; Caryl et al. 2000; Higgins et al. 2005; Ferdous et al. 2012; Chambon et al. 2018). Differences between the paternal and maternal genomes, including sequence duplications, inversions or translocation can lead to non-allelic alignments and may result in loss of genetic information, chromosome mis-segregation, apoptosis and ultimately sterility (Sasaki et al. 2010). While the afore mentioned meiotic failures are associated with rare pathological conditions, genomic loci that are comprised of repetitive sequences have to be maintained in every meiosis and present a potential challenge for DNA repair and recombination systems. For instance, telomeres, centromeres and ribosomal RNA gene are comprised of repetitive sequence elements (Murata et al. 1994; Copenhaver and Pikaard 1996a; Heacock et al. 2004).

Ribosomal RNA genes (or rDNA), encoding the RNA subunits of ribosomes, represent one of the most important and highly conserved repetitive regions in the genomes of eukaryotes (Bell et al. 1989; Shaw and Jordan 1995). They are transcribed by a dedicated set of DNA dependent RNA polymerases forming the Nucleolus Organizer Regions (NORs) and ultimately are part of the nucleolus (Preuss and Pikaard 2007). The nucleolus is a complex sub-nuclear structure and the site of rRNA gene transcription, rRNA processing, and ribosome assembly. Ribosomes represent an essential and evolutionary deeply rooted intersection, translating genetic information from RNA templates into proteins. In higher eukaryotes the ribosomal RNA components are encoded by the 5S rRNA and the 45S rRNA genes (Weis et al. 2015). Although both together ultimately build ribosomes, they differ in complexity and evolutionary dynamics. They are typically physically separated in the genome. The *A. thaliana* (ecotype Col-0) 5S rDNA is located on chromosomes III, IV and V and the 45S rDNA is located in sub-telomeric clusters on chromosomes II and IV (Copenhaver et al. 1995; Copenhaver and Pikaard 1996b; Murata et al. 1997). Such a configuration is wide-spread among higher eukaryotes, for instance in mouse the 45S rDNA loci are located at the sub-telomeric regions of the acrocentric chromosomes (XI, XII, XV, XVI, XVIII, XIX) and the human rDNA loci are located on chromosomes XIII, XIV, XV, XXI and XXII (Gibbons et al. 2015). In Arabidopsis, genome size variations between ecotypes have been attributed to rDNA cluster size variations (Long et al. 2013). In the ecotype Col-0 each of the two 45S rDNA clusters contains about 400 repeats and are 4 Mb in length (Copenhaver and Pikaard 1996b). Loss of rDNA units is a hallmark of aging (McStay 2016) and reduced fitness (Muchova et al. 2015; Warmerdam et al. 2016; Lu et al. 2018). Therefore 45S rDNA repeat numbers need to be maintained in a certain species-specific window, both during the life span of an individual and within a population bridging multiple generations. (Larsen and Stucki 2015; Muchova et al. 2015; McStay 2016; Warmerdam et al. 2016).

Artificially induced DSBs in mammalian cells have shown that breaks within rDNA loci lead to transcriptional shut-down and re-organization of the nucleolus with the formation of so-called nucleolar caps including the damaged rDNA repeats. Within these caps, unscheduled DNA synthesis, presence of specific DNA repair proteins and corresponding functional studies indicate active DNA repair (Larsen and Stucki 2015; Sluis et al. 2015; McStay 2016; Warmerdam et al. 2016). It appears that a first wave of fast repair operates via NHEJ and that persisting damage is repaired via HR. (Harding et al. 2015; Larsen and Stucki 2015; Sluis et al. 2015; McStay 2016). Furthermore, a number of genetic disorders that include compromised DNA repair phenotypes, including Bloom syndrome which is characterized by mutations in the BLM helicase, are associated with loss of rDNA stability and repeat number (Schawalder et al. 2003).

In plants, little is known about the protein factors that are involved in the repair of rDNA. Arabidopsis plants with dysfunctional FAS1, a subunit of the Chromatin assembly factor complex (CAF-1) involved in histone replacement, are enriched in γH2Ax (a histone variant present at DNA DSBs) at the rDNA, show rDNA bridges in mitotic anaphase and suffer from rDNA loss. The rDNA loss phenotype is alleviated in a *rad51*^-/-^ background, indicating that it is a consequence of HR (Muchova et al. 2015; Varas et al. 2017)

DSB formation is an essential part of meiosis. In the yeast *Saccharomyces cerevisiae* meiotic DSBs, and therefore potential non-allelic recombination events, are suppressed within the rDNA array by the histone deacetylase Sir2, the AAA+ ATPases Pch2 and Orc1 (also part of the origin of replication complex) (Gottlieb and Esposito 1989; Ozenberger and Roeder 1991; Mieczkowski et al. 2007; Vader et al. 2011). Sequencing of SPO11-associated oligonucleotides in yeast suggests that very few, if any, meiotic DSBs occur in the rDNA (Pan et al. 2011). A related study in maize estimated that only 0.3% of meiotic DSB sites are located in rDNA loci (He et al. 2017).

Given the importance of rRNA genes and their central function for any living organisms we set out to identify the molecular mechanisms that maintain rDNA clusters over generations in Arabidopsis. Our study demonstrates that both 45S rDNA clusters are transcriptionally active during meiotic prophase and that they associate with the nucleolus. This appears to be a key event to shield the rDNA from acquiring canonical meiotic chromatin modifications, meiotic cohesin and SPO11-dependent DSBs. DNA breaks within the rDNA clusters are repaired in a RAD51-independent, but LIG4-dependent manner, establishing that it is NHEJ and not HR that maintains rDNA integrity during meiosis. Utilizing an ectopically integrated rDNA repeat we validate our findings and suggest that the rDNA constitutes a HR-refractory genome environment.

## Results

### Both 45S rDNA loci are transcribed at the onset of meiosis and localize within the nucleolus

In order to understand how 45S rDNA repeat units are maintained during meiosis we first determined their spatial and temporal distribution and their transcriptional status. Utilizing a FISH probe against the 45S rDNA region and an antibody that binds DNA:RNA hybrids (S9.6, R-loops) (García-Rubio et al. 2015) we established that early in meiosis the rDNA arrays on chromosomes 2 and 4 are transcribed (n=12 cells) (Figure 1A). In contrast, only the NOR on chromosomes 4 (NOR4) in *Arabidopsis* Col-0 adult somatic cells is actively transcribed, while the other NOR (NOR2) is silenced (Mohannath et al. 2016) (n=10 cells) (Figure 1B). We validated these findings by extracting meiocytes from young buds and of different stages and performing rRNA expression analysis of ecotype-specific length polymorphisms (short repetitive sequences) present in the 3′ ETS (External Transcribed Sequence) (Frederic Pontvianne et al. 2007; Durut et al. 2014; Mohannath et al. 2016). In adult leaves only rRNA variants 2 and 3 from NOR4 are expressed, while variants 1, 3 and 4, residing on both NOR2 and NOR4 are detected in early and late meiocytes. In siliques, containing fertilized embryos, all rRNA variants are strongly expressed (Figure 1C).

**Figure 1:**
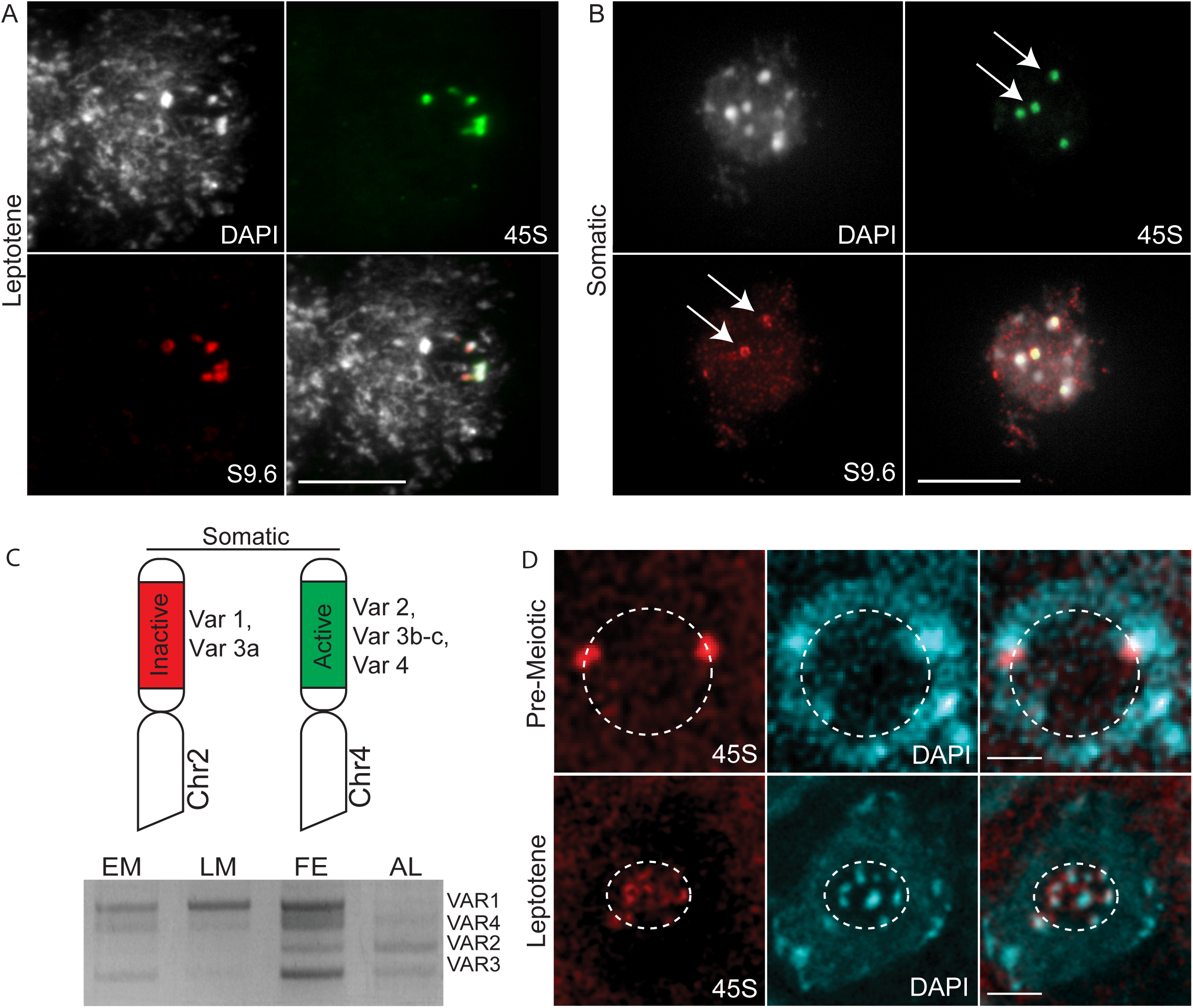
Both NORs are highly dynamic regions and transcribed during meiosis. (A-B) Nuclei stained with S9.6 antibody directed against DNA:RNA hybrids (red), 45S rDNA visualized with a specific FISH probe (green) and DNA stained with DAPI (white). (A) Spread nucleus of a pollen mother cell at leptotene stage. All NORs have a strong S9.6 signal. (B) Spread somatic cell nucleus. White arrow heads indicate the two active NORs (green) colocalizing with the S9.6 signal (red). Size bars = 10 µm. (C) Top: Illustration of *Arabidopsis* chromosomes 2 and 4; the localisation of NORs, the corresponding 45S variants and their transcriptional status in somatic cells is indicated. Bottom: Expression analysis of rDNA variants by RT-PCR during prophase I (early meiosis, EM), post-prophase I and meiosis II (late meiosis, LM), siliques containing fertilized embryos (FE) and in somatic tissue (adult leaves, AL). The agarose gel separates the 4 DNA bands representing rDNA 3’ETS variant 1 (VAR1), 2 (VAR2), 3 (VAR3) and 4 (VAR4). (D) Single optical layer of meiotic nuclei after a whole mount FISH preparation. 45S rDNA has been visualized via FISH using a specific probe (red) and DNA has been stained with DAPI (cyan). Dashed circles highlight the nucleolus. Size bars = 2 µm

To examine whether these transcriptional states are associated with larger scale chromosome dynamics we performed whole mount FISH on anthers to generate three-dimensional reconstructions of meiotic cells. After pre-meiotic S-phase, the NORs are localized in a canonical somatic configuration around the nucleolus (Pontvianne et al. 2013) (n=30 cells from different anthers), while during meiotic prophase, from leptotene onwards, they localize within the nucleolus (n=30 cells from different anthers) (Figure 1D; Videos 1, 2). Moreover, from zygotene onwards, both NORs form a unified structure and only disengage at the end of meiotic prophase I, during diakinesis, when the nuclear envelope breaks down and paired chromosomes condense in preparation for segregation. In agreement with the rRNA expression data, DNA:RNA hybrids, which mark actively transcribed genes, colocalized with the rDNA throughout prophase I of meiosis (Suppl. Figure S1). We conclude that both NORs become actively transcribed at the onset of meiosis and that this is correlated with their localization in the nucleolus.

### Meiotic rDNA is embedded in a unique chromatin environment

With the rDNA loci residing in the nucleolus from leptotene onwards they are partitioned from the rest of the chromatin during meiosis. To probe the functional relevance of this sequestration we analysed potential differences in chromatin architecture and modification. During leptotene and zygotene, antibodies directed against the axis protein ASY1 or the SC protein ZYP1 fail to co-localize with the 45S signal (rDNA) (n=17) (Figure 2A). In pachytene, the formation of the SC corresponds with extended stretches of ZYP1 along the paired chromosomes, and the depletion of ASY1 (Higgins et al. 2005). Remarkably, at this stage the rDNA loci acquire a prominent ASY1 signal, while the rest of the chromatin is largely devoid of it (n=25) (Figure 2A). Whole mount immuno-FISH, which preserves the spatial relation of nucleolus and chromatin within the nucleus, using probes for rDNA (45S) and an antibody marking the chromatin axis (ASY1) revealed that the nucleolus itself is free of ASY1 (n=32 cells) (Figure 2B; Videos 3, 4). To understand the 3D relationship of rDNA, axis and SC we simultaneously stained for ASY1 and ZYP1 on spread chromatin of PMCs at pachytene, and imaged the meiocytes using super-resolution confocal microscopy. At a 160 µm resolution it is apparent that the strong ASY1 signals represent four spatially separated chromatin stretches devoid of any ZYP1 signal. The previous experiments established that these stretches represent the NOR regions of chromosome 2 and 4. Taken together, this demonstrates that in contrast with the rest of the genome, the homologous chromosomes in the NOR regions do not undergo synapsis (n=5 cells) (Figure 2C, Video 5).

**Figure 2:**
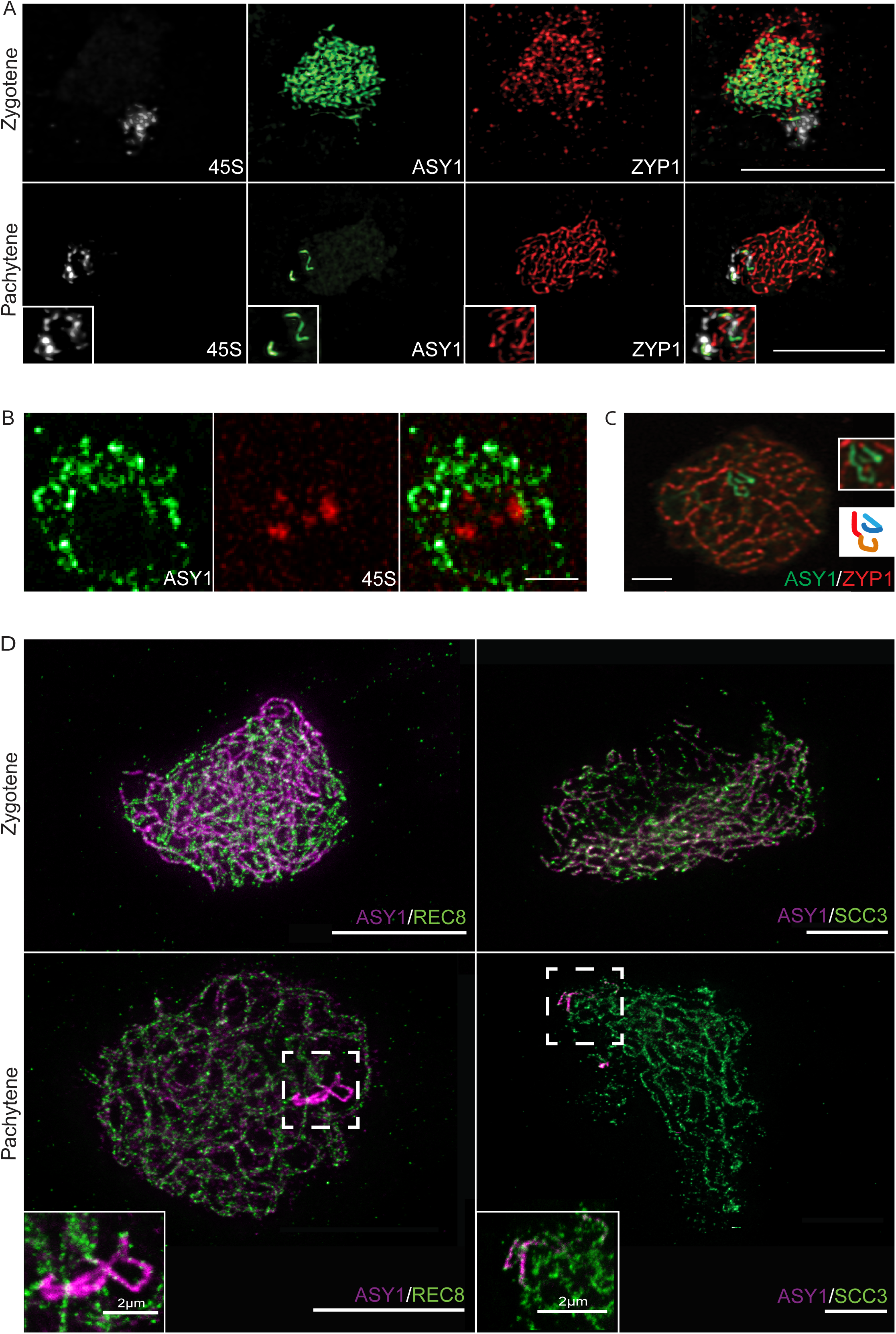
The rDNA acquires distinct chromatin characteristics during meiosis. (A) Immuno-FISH spreads of PMCs at zygotene and pachytene stages. The meiotic axis was stained with an anti-ASY1 antibody (green), the SC with anti-ZYP1 antibody (red) and the DNA was hybridized with a 45S rDNA probe (white). Scale bars = 10 µm. (B) Single optical layer (160 nm) of a meiotic nucleus after whole mount immuno-FISH preparation. 45S rDNA has been visualized via FISH using a specific probe (red) and the meiotic axis with a specific antibody directed against the protein ASY1 (green). Size bar = 2 µm. (C) Single optical layer (160 nm) of a spread meiotic nucleus at pachytene stage stained for the axis with anti-ASY1 antibody (green) and for the SC with anti-ZYP1 antibody (red). The image has been acquired with an AiryScan super-resolution unit on a confocal microscope. Size bar = 2 µm. The cartoon highlights the chromatin stretches colocalizing with the NORs during pachytene stage and depicts the two separated homologues of chromosomes 2 and 4. (D) Super-resolution images of spread meiotic nuclei at zygotene and pachytene stages stained for the axis with an anti-ASY1 antibody (magenta) and for cohesin subunits with anti-REC8 or anti-SCC3 antibodies (green). The nuclei were imaged with an Abberior STEDYCON microscope at 40-60 nm resolution. The boxed areas have been enlarged to highlight the localization of cohesin subunits relative to the axis protein ASY1. Size bars = 10 µm.

To investigate the molecular basis of the chromosome morphologies we observed, we used immuno-localization of antibodies that recognize the meiosis specific cohesin subunit REC8 (Cai et al. 2003) and the cohesin complex partner SCC3 (Chelysheva et al. 2005). We used super-resolution microscopy and found that both REC8 and SCC3 localize to chromosome axes during zygotene (n=18 cells, REC8; n=11 cells, SCC3). During pachytene, with ASY1 predominantly localizing to rDNA, only SCC3 but not REC8 co-localized with ASY1 (n=7 cells, REC8; n=8 cells, SCC3) (Figure 2D, Suppl. Figure 2). This demonstrates that rDNA acquires a cohesin complex that excludes the canonical, meiosis-specific REC8 kleisin subunit (Cai et al. 2003).

Several chromatin modifications are enriched in the 45S rDNA arrays. Mono-methylation of H3K27 is strongly enriched at the somatically silenced NOR2 and is introduced by the two methyltransferases ATXR5 and ATXR6 (Mohannath et al. 2016). In contrast, H4Ac4, a known euchromatin mark, is mainly associated with the active NOR4 in adult somatic cells. Histone Deacetylase 6 (HDA6) has been shown to remove H4 acetylation from NOR2, contributing to its silenced state (Probst et al. 2004; Earley et al. 2006; To et al. 2011). We performed immuno-FISH on spread chromatin from PMCs and found that in zygotene, when all 45S rDNA resides within the nucleolus and is assembled into a unified structure, both NORs are enriched in H3K27me1 but depleted in H4Ac4 (n=12 cells, H3K27me1; n=16 cells, H4Ac) (Figure 3, Suppl. Figure 3), indicating that their somatically distinct chromatin environments have been matched, and that at this stage both NORs carry marks of a repressed chromatin state. To validate these results we performed similar analyses in the corresponding mutant backgrounds. The *atxr5 atxr6* double mutants do not show H3K27me1 marks on meiotic NORs (Suppl. Figure 3A) and meiocytes of the *hda6-6* mutant display NORs that are enriched in H4Ac4 (Suppl. Figure 3B). Moreover, the NORs in *hda6-6* mutants do not form a compact, unified structure, with most cells displaying two separate rDNA clusters (90% of all observed cells at zygotene/pachytene stage; n=31) indicating that H4 deacetylation is needed to maintain rDNA clustering within the nucleolus during meiosis. Similarly, in *atxr5 atxr6* double mutants 30% of all observed cells (n=26) have separated rDNA clusters (Figure 4A, Suppl. Figure 4A). Taken together, these results suggest that the 45S rDNA arrays are sequestered in the nucleolus during prophase I of meiosis and embedded in a unique chromatin environment.

**Figure 3:**
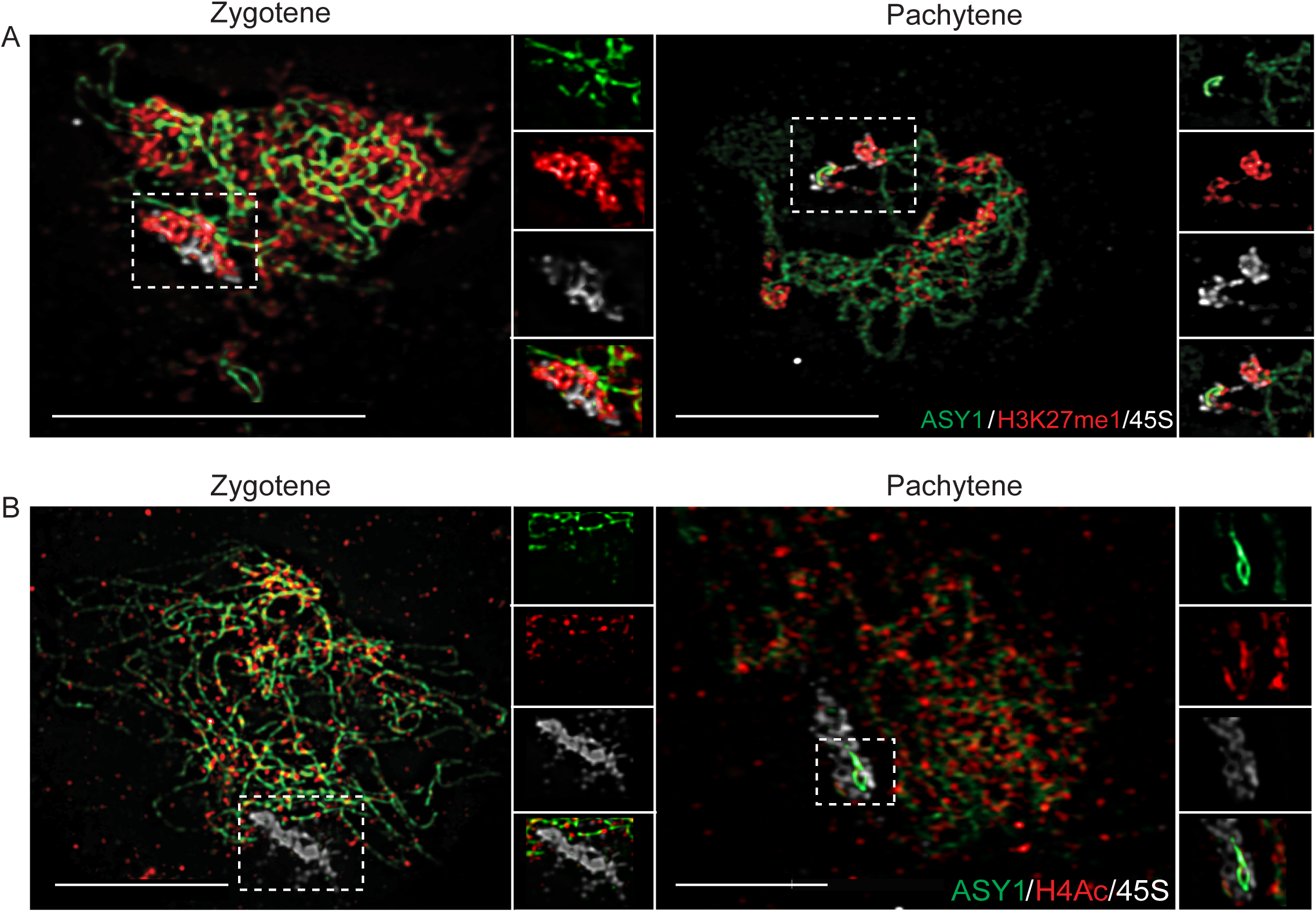
The rDNA acquires a specific chromatin environment during meiosis. Immuno-FISH spreads of meiotic cells at zygotene/pachytene stages. The axis has been stained with an anti-ASY1 antibody (green) and the histone modification H3K27me1 with a specific antibody (red). The 45S rDNA has been visualized with a specific FISH probe (white). Immuno-FISH spreads of meiotic cells at zygotene/pachytene stages. The axis has been stained with an anti-ASY1 antibody (green) and the histone modification H4Ac with a specific antibody (red). The 45S rDNA with has been visualized with a specific FISH probe (white). For the boxed areas individual channels are shown to highlight the localization of the histone modifications relative to the axis protein ASY1. Size bars = 10 µm.

**Figure 4:**
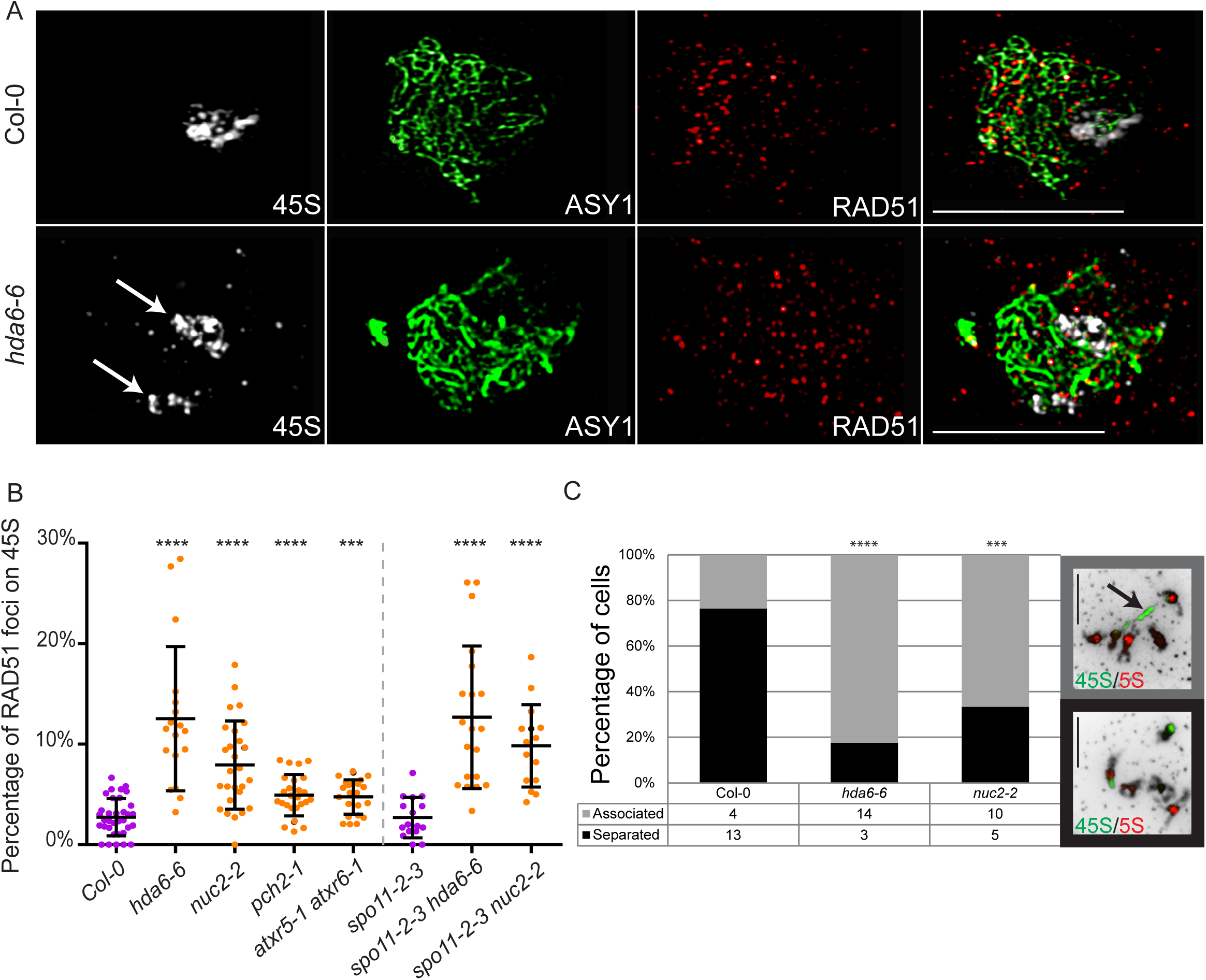
The nucleolus shields rDNA from meiotic DSB formation and deleterious HR. (A) Immuno-FISH spreads of wild-type (Col-0) and mutant (*hda6-6*) PMCs at zygotene. The axis has been stained with an anti-ASY1 antibody (green) and the recombinase RAD51 with a specific antibody (red). The 45S rDNA has been visualized with a specific FISH probe (white). White arrows indicate separated NORs in the *hda6-6* mutant. (B) Percent RAD51 foci colocalizing with the 45S rDNA probe in relation to the total number of RAD51 foci per nucleus. Statistical analysis was performed by using a Mann Whitney test. (C) Graph depicting the percentage of nuclei with associated or clearly separated NORs counted from PMC spreads at diakinesis stage. The DNA has been hybridized with a 45S rDNA probe (green) and a 5S rDNA probe (red). Statistical analysis was performed by using a binary logistic regression. Arrow indicates two fused NORs. Size bars = 10 µm.

### The nucleolus shields rDNA from SPO11-mediated DSB formation and homologous recombination

To determine whether the meiotic chromatin environment of the 45S rDNA influences DSB formation and repair we used immuno-FISH with a RAD51 antibody (Kurzbauer et al. 2012) and a 45S DNA probe on chromatin spreads of PMCs to monitor HR progression (Figure 4A, Suppl. Figure 4A). On average only 4.39 ± 4.04 (2.7%) of all RAD51 foci were localized at the rDNA (166 ± 65 RAD51 foci at leptotene/zygotene in total; n=29). This is significantly lower (p=0.0001) than the expected 11.8 RAD51 foci on rDNA, assuming a random distribution across the genome (on average 7.6% of the area of a spread meiotic nucleus is rDNA; n=15; in agreement, 7% of the *A. thaliana* Col-0 genome has been estimated to be rDNA (Copenhaver and Pikaard 1996a)). This indicates a mechanism that shields the rDNA from acquiring RAD51 foci. To investigate which protein factors may be facilitating this safeguarding mechanism we evaluated RAD51 foci co-localizing with the rDNA in various mutant backgrounds. Having established previously that HDA6 and ATXR5/6 are instrumental for a specific rDNA chromatin environment and also for nucleolus morphology in meiosis (see above) (Earley et al. 2006; Pontvianne et al. 2012) we investigated RAD51 foci numbers in spreads of PMCs of *hda6-6* plants and observed a significant increase in RAD51 foci co-localizing with the rDNA compared to wild type (12.5% on rDNA; n=18; p<0.0001) (Figure 4A-B, Suppl. Fig. 4A). We also analysed plants deficient in the conserved plant nucleolin gene NUC2 that have altered rDNA transcription and NOR morphology compared to wild type plants (Frederic Pontvianne et al. 2007; Durut et al. 2014). In *nuc2-2* mutants significantly elevated numbers of RAD51 foci co-localize with the rDNA compared to the wild type (7.9% on rDNA; n=27; p<0.0001) (Figure 4B, Suppl. Fig. 4C). We also investigated *pch2* mutant lines (Lambing et al. 2015) motivated by the findings in yeast that the PCH2 protein is instrumental in protecting the repetitive rDNA from recombination events (Vader et al. 2011) and detected a significant increase of RAD51 foci numbers on rDNA compared to wild type (4.9% on rDNA; n=26; p<0.0001). Also, in *atxr5 atxr6* double mutants a significant increase of RAD51 foci numbers co-localizing with the rDNA was observed (4.7% on rDNA; n=23; p<0.001) (Figure 4B, Suppl. Figure 4C). It is important to note that the total (genome-wide) number of RAD51 foci in all tested mutant lines did not significantly differ from wild type (or *spo11-2-3*) (p>0,39), apart from plants carrying the *nuc2-2* mutant allele with slightly reduced overall RAD51 foci numbers (p<0.05) (Suppl. Figure 4B).

To test if the increase of 45S rDNA-associated RAD51 foci in *hda6-6* and *nuc2-2* mutants is caused by an increase of SPO11-dependent DSBs we generated *spo11-2-3 hda6-6* and *spo11-2-3 nuc2-2* double mutants. We observed a few rDNA-associated RAD51 foci in *spo11-2-3* mutants (2.7% on rDNA; n=16). In *spo11-2-3 hda6-6* double mutants however we noted a significant increase of RAD51 foci co-localizing with the rDNA when compared to *spo11-2-3* single mutants (12.6% on rDNA; n=18; p<0.0001). We conclude that the elevated number of 45S rDNA-associated RAD51 foci in *hda6-6* mutants are not SPO11-dependant. The increase of 45S rDNA-associated RAD51 foci observed in *nuc2-2* mutants also appears independent of SPO11. In *spo11-2-3 nuc2-2* double mutants the number of RAD51 foci co-localizing with the rDNA is significantly increased compared to *spo11-2-3* single mutants (9.8% on rDNA; n=15; p<0.0001) (Figure 4B, Suppl. Figure 4B).

These results demonstrate that in wild type plants the rDNA, residing within the nucleolus during most of meiotic prophase I, is protected from canonical, SPO11-mediated DSB formation and also from the homologous recombination protein RAD51. Based on this, we hypothesize that the elevated number of 45S rDNA-associated RAD51 foci in *hda6-6* and *nuc2-2* meiocytes may represent locations of homologous recombination DNA repair, with the potential of non-allelic interactions among rDNA repeats.

To test this hypothesis, we performed FISH using probes for 45S and 5S rDNA loci, allowing us to distinguish the 5 Arabidopsis chromosomes. In wild type plants only 24% (n=17) of meiocytes at diakinesis show interconnected 45S rDNA signals from chromosomes 2 and 4, but in *hda6-6* and *nuc2-2* 82% (n=17; p=0.0004) and 59% (n=15; p=0.015) of meiocytes respectively had these connections (Figure 4C).

Taken together, the data suggest that both 45S rDNA loci are sequestered into the nucleolus during meiotic prophase I where they are protected from SPO11 mediated DSB formation, shielded from RAD51 loading and constrained to prevent interactions that have the potential for non-allelic recombination. Nevertheless, some DNA breaks occur in the 45S rDNA, as inferred by the limited loading of RAD51. In mutant backgrounds that do not preserve the nucleolus, these breaks may preferentially load RAD51 and be repaired via HR. The breaks may result from collisions between replication and transcription machineries (Aguilera and Gaillard 2014).

### Repair of rDNA depends on NHEJ mechanisms

In order to directly investigate break formation in the 45S rDNA and identify the corresponding DNA repair mechanism we quantified rDNA integrity at the end of meiosis in wild type and mutant lines. We performed FISH using 45S rDNA probes on meiocytes at anaphase II to tetrad stage and counted the number of individual 45S rDNA signals. Eight non-fragmented rDNA signals are expected in each wild type tetrad (2 NORs per haploid microspore cell). 45S rDNA fragmentation are expected to result in tetrads with more than 8 signals. We observed no more than 8 45S rDNA signals in wild type meiocytes (n=25), but saw significantly more tetrads with greater than 8 fragments in *lig4-4* (21%, n=32, p=0.0029 compared to Col-0) and *mre11-4* (16%, n=31, p=0.017) mutants (Figure 5A – C, Suppl. Figure 5). LIGASE4 is a well-conserved hallmark factor in the canonical non-homologous end joining (C-NHEJ) DNA repair pathway (Friesner and Britt 2003). MRE11 required for both HR and micro-homology mediated end joining (MMEJ), an alternative NHEJ pathway. To test if HR has a role in DNA repair of 45S rDNA during meiosis we quantified 45S rDNA FISH signals in late meiosis II meiocytes in *com1-1* and *rad51-1* mutants. SPO11 mediated breaks are not repaired in these two mutants, leading to severe chromosome fragmentation during meiosis (Li et al. 2004; Uanschou et al. 2007), but the 45S rDNA is not fragmented (Figure 5C, Suppl. Figure 5). Consistent with our earlier observations that SPO11 is not mediating DNA break formation in 45S rDNA (Figure 4) (Pan et al. 2011; He et al. 2017), the fragmentation in *lig4-4* and *mre11-4* mutants is not alleviated in the corresponding *spo11-2-3 lig4-4* (22% of cells have > 8 rDNA signals, n=22, p=0.004) and *spo11-2-3 mre11-4* (14% of cells have > 8 rDNA signals, n=27, p=0.0185) double mutants (Figure 5C, Suppl. Figure 5).

**Figure 5:**
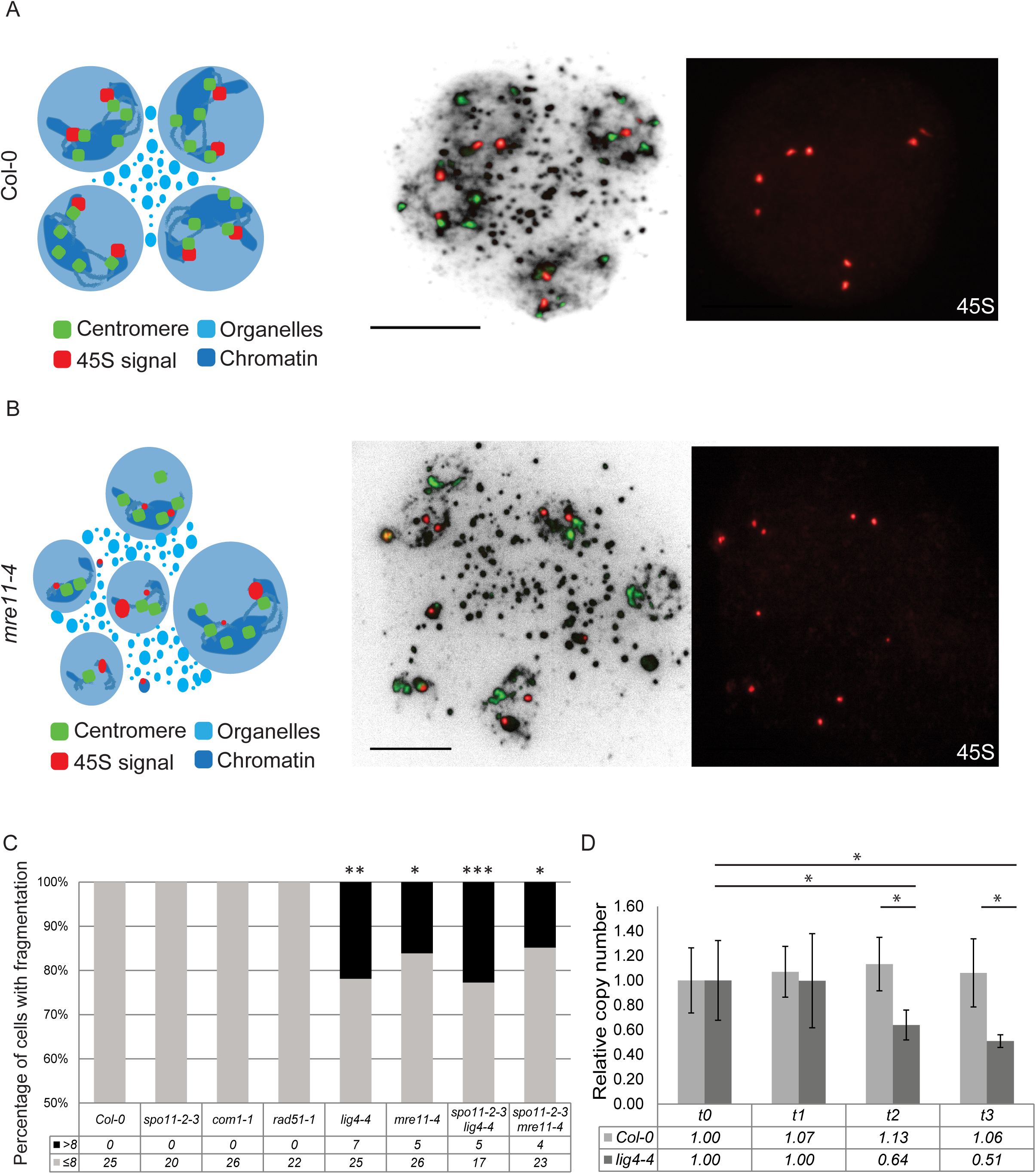
The rDNA is repaired by NHEJ. (A) Left: cartoon representing a meiotic tetrad stage of a wild type (Col-0) meiocyte. Middle: spread nuclei of a meiocyte at tetrad stage followed by FISH to visualize the 45S rDNA (red) and the centromeres (green). Right: Panel depicts only the FISH signals that visualize the 45S rDNA (red), showing eight distinct signals. (B) Left: cartoon representing a meiotic tetrad-like stage of a *mre11-4* mutant meiocyte. Middle: spread nuclei of a meiocyte at tetrad-like stage followed by FISH to visualize the 45S rDNA (red) and the centromeres (green). Right: Panel depicts only the FISH signals that visualize the 45S rDNA (red), showing ten distinct signals. (C) Quantification of 45S signals per individual meiocyte. Only stages from anaphase II to tetrad were analysed. *lig4-4, mre11-4, spo11-2-3 lig4-4* and *spo11-2-3 mre11-4* show an increase in 45S foci numbers compared to Col-0. Statistical analysis was performed by using a binary logistic regression. (D) Graph depicting the loss of 45S copy numbers in a *lig4-4* mutant plant, compared to wild type (Col-0), grown under short day conditions. t0 to t3 refers to the age (young to old) of the meristem when leaves were generated. Statistical analysis was performed by using a t-test. Size bars = 10 µm.

These results provide further evidence that meiotic DSBs in the NORs are not generated by SPO11. As mentioned above, they may be a by-product of collisions between replication and transcription machineries. Furthermore, it appears that DSB repair in the 45S rDNA during meiotic prophase I is: independent of COM1 and RAD51, which are used in HR but not C-NHEJ or MMEJ; dependent on LIG4, which is used in C-NHEJ but not MMEJ or HR; and dependent on MRE11, known for its role in HR, C-NHEJ and MMEJ (McVey and Lee 2008).

To test whether NHEJ is also essential for repair of 45S rDNA-associated DSBs in somatic cells we determined the relative rDNA copy number by qPCR in wild type and *lig4-4* mutant lines grown side-by-side. We picked rosette leaves of different sizes from plants grown for 67 days under short-day conditions which strongly delays the transition from a vegetative meristem, producing leaves, to a generative meristem producing flowers. This allowed us to assay cumulative 45S rDNA phenotypes within meristematic tissues. Newer leaves (from older meristems) had a significant reduction in 45S rDNA copy number in *lig4-4* plants compared to older leaves that had been produced earlier during meristematic growth (Figure 5D), but wild type plants showed no differences across the time course (p≤0.03). The effect is most pronounced in young leaves from relatively old meristems. A strong candidate for the loss of 45S rDNA copies is the inability to repair DSBs with pathways typically utilized in wild type backgrounds and instead repair them with deletion prone pathways. This suggests a role for LIG4 and C-NHEJ in the repair of 45S rDNA lesions in wild type dividing, somatic cells.

### The rDNA creates an HR-refractory chromatin environment

To test the idea that the 45S rDNA is sequestered in an HR-refractory chromatin environment during meiosis, we generated plant lines carrying single ectopic rDNA insertions, each containing only a few rDNA units, to test the effect of rDNA on meiotic recombination (Figure 6, Suppl. Figure 6). These experiments also serve as a control for any unrecognized genetic element at the native NORs. The transformation (T-DNA) vector we used contains one complete rDNA unit (variant 1), and includes a unique sequence of 20 nucleotides within a variable portion of the 25S rDNA region, enabling its specific detection (Wanzenböck et al. 1997). Only plant lines with a 3:1 segregation pattern, indicating a single locus insertion, were used. The insertion sites were mapped, ectopic rDNA copy numbers evaluated and active transcription confirmed (ErDNA2, Ectopic rDNA on chromosome 2, six copies integrated at nucleotide position 1,704,488; ErDNA3, Ectopic rDNA on chromosome 3, one copy integrated at nucleotide position 10,863,135; ErDNA5, Ectopic rDNA on chromosome 5, two copies integrated at nucleotide position 4,999,535) (Suppl. Figure 6A, B). To control for T-DNA related effects, not functionally linked to the rDNA, we obtained T-DNA insertion lines from T-DNA insertion collections nearby the genomic positions of ErDNA3 and ErDNA5 (TDNA3: SALK_137758; TDNA5: SAIL_713_A12). We generated a negative control using CRISPR/CAS9 (Fauser et al. 2014; Steinert et al. 2015) to delete the rDNA portion, but leaving the T-DNA backbone intact, within the ErDNA5 transgene (ErDNA5-del) (Suppl. Figure 6C).

**Figure 6:**
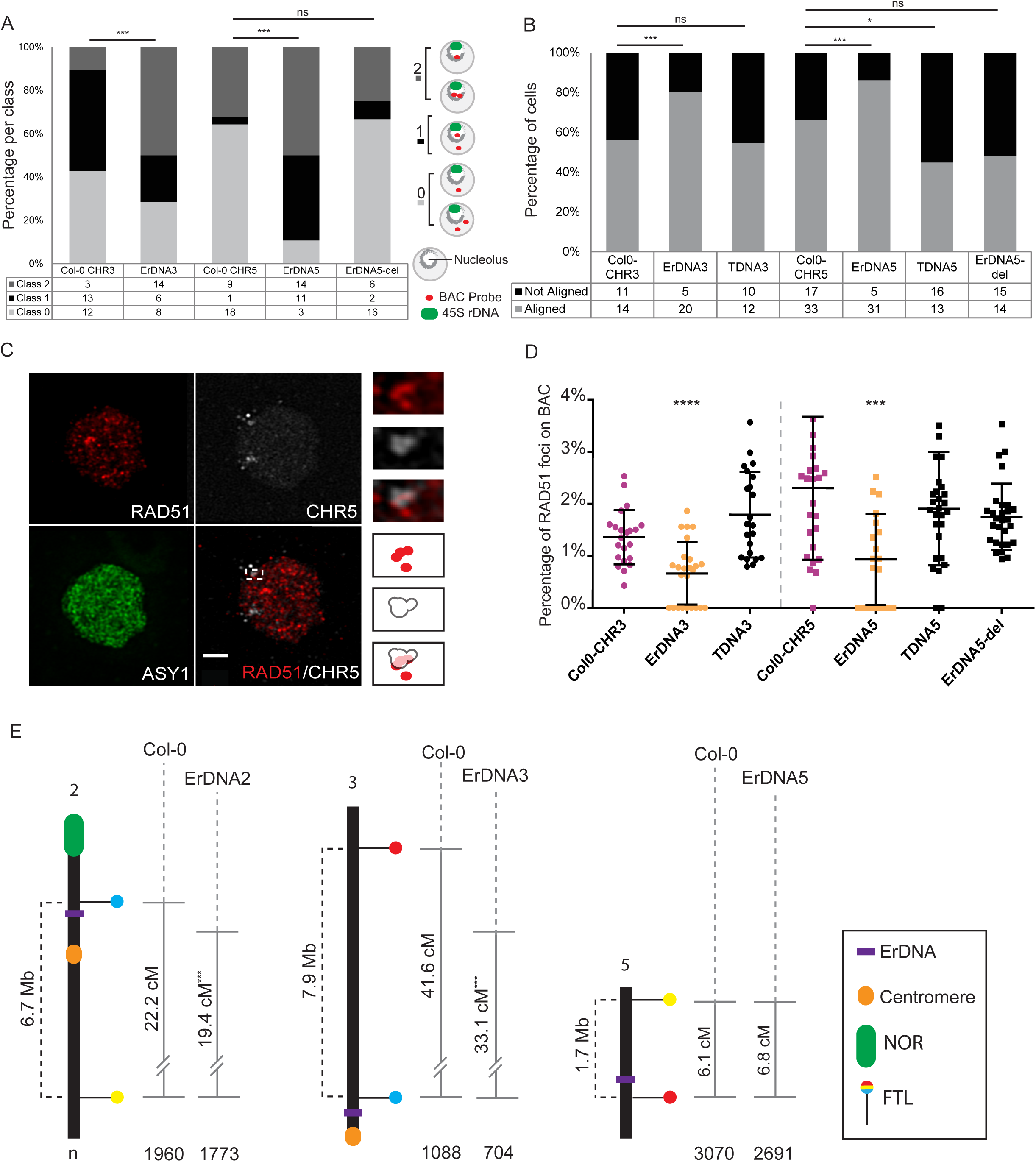
Ectopically integrated rDNA units associate with the nucleolus, promote homologue pairing and suppress meiotic recombination. (A) Graph depicts incidence of different classes of nucleolus association of genomic regions with (ErDNA3, ErDNA5) or without (Col-0) the ectopically integrated rDNA and in a control line (ErDNA5-del). Data has been obtained from whole mount FISH preparations of anthers. DNA was hybridized with BAC probes for the 500kb regions of interest and with a 45S rDNA probe. Statistical analysis was performed by using a binary logistic regression. (B) Graph depicts incidence of homologous alignments of genomic regions with (ErDNA3, ErDNA5) or without (Col-0) the ectopically integrated rDNA and in control lines (T-DNA, ErDNA5-del). Data has been obtained from Immuno-FISH preparations. DNA was hybridized with BAC probes for the 500kb regions of interest. Statistical analysis was performed by using a binary logistic regression. (C) Immuno-FISH spreads of wild type (Col-0) PMC at zygotene (single 160nm optical layer shown). The axis has been stained with an anti-ASY1 antibody (green) and the recombinase RAD51 with a specific antibody (red). The 500kb region of interest on chromosome 5 has been visualized with a specific FISH probe (white). Small panels on the right demonstrate RAD51 – FISH probe colocalization analysis. Only RAD51 foci that had an overlap of at least 50% with the FISH signal were counted as “colocalizing”. (D) Graph showing the percentage of RAD51 foci colocalizing with a 500kb region on chromosome 3 or chromosome 5 in relation to the total RAD51 foci counts per meiocyte. Colocalizing events were counted in wild-type (Col-0), ErDNA lines and control lines (T-DNA, ErDNA5-del). Statistical analysis was performed by using a Mann Whitney test. (E) Crossover frequencies measured by fluorescent pollen markers (FTLs) in three different intervals. Genetic distances were analysed in the absence (wild type, Col-0) or presence (ErDNA2, ErDNA3 and ErDNA5) of ectopic rDNA repeats. Statistical analysis was performed by using a binary logistic regression. Size bar = 2 µm.

We designed FISH probes that hybridize to a 500kb window surrounding the ectopic rDNA integration sites on chromosomes 3 and 5. We performed whole-mount immuno-FISH experiments to determine the relative position of these regions in relation to the nucleolus, in wild type plants compared to ErDNA3, ErDNA5 and ErDNA5-del (Figure 6A, Suppl. Video 6). The presence of ectopic rDNA caused the corresponding 500 kb regions on chromosome 3 or 5 to localize or associate with the nucleolus. To quantify the localization patterns we sorted the FISH signals into 3 classes: Class 2 has signals (fused or not) associated with or residing in the nucleolus; Class 1 has one signal, associated with or residing in the nucleolus and one not; Class 0 has signals (fused or not), that are not associated with and do not reside within the nucleolus. In wild type plants the chromosome 3 integration site is found in class 2 configurations in 10%, in class 1 in 46% and in class 0 in 44% of cells (n=28). In contrast, in ErDNA3 lines with a single rDNA unit in the same 500kb window, the signals are in class 2 configurations in 50%, in class 1 in 21% and in class 0 in 29% of cells (p=0.01, n=28). Similarly, in wild type plants the integration site on chromosome 5 is found in class 2 configurations in 32%, in class 1 in 3% and in class 0 in 65% of cells (n=28). In contrast, ErDNA5 lines, with two rDNA units in the same 500kb chromosome 5 window, had signals in class 2 configurations in 50%, in class 1 in 39% and in class 0 in 11% of cells (p=0.001, n=28). The signal configurations in the ErDNA5-del control line lacking ectopic rDNA, resembles wild type with class 2 configurations in 25 %, class 1 in 8% and in class 0 in 67% of cells (p=0.45, n=24) (Figure 6A). Furthermore, the genomic regions on chromosomes 3 and 5 were significantly more often paired with their corresponding homologous partner sites when the ectopic rDNA was present. In the ErDNA3 line the region on chromosome 3 was paired in 79% of cells in leptotene/zygotene stage (n=25; p=0.035), while in wild type and TDNA3 control cells only 58% (n=24) and 56% (n=22; p=0.46) had paired homologous loci respectively. Similar results were obtained also for ErDNA5 with 86% of cells (n=36; p=0.01) having paired homologous loci in the presence of ectopic rDNA while in wild type only 66% were paired (n=50). In the TDNA5 and ErDNA5-del control lines only 45% (n=29; p=0.03) and 48% (n=29; p=0.07) of cells had paired loci respectively (Figure 6B).

These observations demonstrate that ectopic rDNA units mirror the characteristics of endogenous NORs. We therefore asked whether the presence of an ectopic rDNA unit also decreases the number of RAD51 foci in its vicinity. We performed immuno-FISH experiments, simultaneously detecting the ErDNA integration site (a 500kb window around the target sites), ASY1 to mark the chromosomal axis (for staging), and RAD51 (Figure 6C). We found that the number of RAD51 foci within the 500kb window is significantly reduced in the presence of the ectopic rDNA in ErDNA5 (0.9% of all RAD51 foci co-localize with the FISH signal, n=19, p=0.0004) compared to wild type (2.3% co-localization, n=26). The TDNA5 (1.9% co-localization, n=28, p=0.7) and ErDNA5-del (1.6% co-localization, n=29, p=0.8) control lines were not significantly different from wild type. We obtained similar results for ErDNA3 with only 0.6% of RAD51 foci co-localizing with the FISH signal (n=25; p≤0.0001), while in wild type and TDNA3 control cells 1.3% (n=24) and 1.7% (n=24; p=0.07) of RAD51 foci colocalized with the FISH signal respectively (Figure 6D). The total number of RAD51 foci were not significantly different in any of the lines (p≥0.7 for all cases) (Suppl. Figure 6D). These results demonstrate that ectopic 45S rDNA units inhibit recombination intermediates in their vicinity, in line with our observations with the endogenous NORs.

To measure the effect of 45S rDNA on crossover frequency directly we measured genetic distances in the presence or absence of ectopic rDNA units. We crossed ErDNA2 with lines carrying flanking transgenic markers in a *qrt1* background. The transgene markers, called FTLs, encode fluorescent proteins expressed by the *LAT52* post-meiotic, pollen specific promoter (Francis et al. 2007).

The qrt1 background causes the pollen products of each meiosis to be released as tetrad thus enabling tetrad analysis. The genetic distance between pairs of FTL marker can be calculated by scoring their segregation in the pollen tetrads (Berchowitz and Copenhaver 2008). We flanked the ectopic 45S rDNA in ErDNA2 at nucleotide position 1,704,488 on chromosome 2 with FTL 1431 (at nucleotide position 1,521,041) and FTL 2269 (at nucleotide position 8,276,753). In wild type plants this 6.7 Mb interval has a genetic distance of 22.2 cM (n=1773 tetrads, 4 plants), in the presence of ectopic 45S rDNA (line ErDNA2-FTL) the genetic distance is significantly reduced to 19.4 cM (n=1688 tetrads, 5 plants; p=0.003) (Figure 6E). We obtained a similar result with ErDNA3 by flanking the ectopic 45S rDNA unit chromosome 3 at nucleotide position 10,863,135 with FTL 1019 (at nucleotide position 1,517,290) and FTL 1046 (at nucleotide position 9,458,743). In wild type plants this 7.9 Mb interval has a genetic distance of 41.6 cM (n=1088 tetrads, 6 plants), while in the presence of ectopic rDNA (line ErDNA3-FTL) the genetic distance is significantly reduced to 31.1 cM (n=730 tetrads, 6 plants; p=0.0009) (Figure 6E). The integrations sites of the ectopic 45S rDNA units in ErDNA2 and ErDNA3 reside within recombination proficient genomic regions (Choi et al. 2013), facilitating detection of changes in recombination frequencies. In contrast, the integration site in ErDNA5 at nucleotide position 4,999,535 on chromosome 5 resides within a genomic region that also contains a recombination deficient “cold spot”. We flanked ErDNA5 with FTL 1143 at nucleotide position 3,760,756 and FTL 2450 at nucleotide position 5,497,513. In wild type plants this 1.7 Mb interval has a genetic distance of 6.1 cM (n=4682 tetrads, 6 plants) and in the presence of ectopic 45S rDNA (line ErDNA5-FTL) the genetic distance is not significantly different (6.8 cM, n=2691 tetrads, 4 plants, p=0.07). These results suggest that CO suppression by 45S rDNA acts locally and in recombination-competent contexts (Choi et al. 2013). The results with ErDNA5 suggest that rDNA CO suppression is not additive with the influence of existing cold spots.

## Discussion

One of the most interesting questions in biology is how genetic information is passed from one generation to the next. What mechanisms mediate the balance between preservation of existing allele combinations and recombination to produce novel unions? Over the last decades many molecular factors had been identified and molecular mechanisms of meiotic recombination elucidated (Hunter 2015; Mercier et al. 2015). However, the question of how repetitive genetic elements are inherited remains open. Sequence repeats create a liability during recombination because they can undergo non-allelic exchange and are a potential source of deletions, duplications, inversions or translocations. Until now, meiotic DNA repair thought to primarily depend on HR. The role of other DNA repair pathways in meiosis, like C-NHEJ and MMEJ, is much more poorly understood and generally only associated with pathological conditions when HR is impaired (Lemmens et al. 2013; Girard et al. 2018). Our study, utilizing the model plant *Arabidopsis thaliana*, supports a model in which the highly repetitive 45S rDNA arrays are protected from SPO11-induced meiotic DSBs as well as the meiotic recombination machinery by their recruitment into the nucleolus and that the SPO11-independent DNA lesions that occur in these domains are repaired via C-NHEJ.

### During meiosis the rDNA creates a unique chromatin environment

Our study demonstrates that in *A. thaliana* Col-0 both 45S rDNA arrays, located on chromosomes 2 and 4, become transcriptionally active at the onset of meiosis in contrast to adult somatic tissues, in which only the chromosome 4 array is active (Tucker et al. 2010). Earlier studies demonstrated that in embryos and seedlings both rDNA arrays are transcriptionally active, until one array is silenced (Frederic Pontvianne et al. 2007). Our data reveals that the initial re-activation occurs before fertilization during meiosis. We detect rRNA variants from both arrays and RNA/DNA hybrids at both 45S regions throughout meiosis, yet it is not clear if transcription is maintained over the entire course of meiosis. In somatic cells, rDNA transcription is silenced upon mitotic entry when the nucleolus disassembles (Klein et al. 1999b). We propose that transcriptional activity of both 45S rDNA arrays is crucial for their subsequent association with the nucleolus in prophase of meiosis I. Consistently, actively transcribed, ectopic rDNA units are also preferentially associated with the nucleolus. Furthermore, Grob *et al.* (Grob et al. 2014) showed that in human HT1080 cells a synthetic NOR sequence would only form a new nucleolus if actively transcribed.

Nucleolar association of 45S rDNA repeats appears to be required to shield them from acquiring canonical meiotic chromatin modifications, meiotic cohesin and SPO11-dependent DSBs. Our analysis of chromatin modifications that had been shown previously to characterize silenced (NOR2, H3K27me1) and active (NOR4, H4Ac) rDNA clusters shows that during meiosis both NOR2 and NOR4 obtain identical patterns. This is consistent with the observation that both arrays are transcribed and associate with the nucleolus. Furthermore, both NORs fuse into a unified structure during meiotic prophase I, indicating that the somatic distinction of the two is alleviated at this stage. An important aspect that sets 45S rDNA apart from the rest of the genome during meiosis I is the exclusion of the axis protein ASY1 during leptotene and zygotene. ASY1 is also involved in establishing inter-homologue bias during meiotic HR (Sanchez-Moran et al. 2007). Based on those observations we hypothesize that its exclusion from the 45S rDNA (and the nucleolus in general) relaxes inter-homologue bias. Only after genome-wide DSB formation and repair has been accomplished, in pachytene, ASY1 localizes to the 45S rDNA, while being removed from other genomic sites. During pachytene the synaptonemal complex (SC) is established between homologous chromosomes, but it is omitted in the region of 45S rDNA. The SC not only stabilizes inter-homologue pairing events, it also promotes the formation of cross-overs (Mercier et al. 2015). Further evidence that the 45S rDNA does not acquire a canonical meiotic chromosome morphology is the absence of the meiosis-specific cohesin subunit REC8 which is part of the meiotic axis and helps promote inter-homologue interactions (Schwacha and Kleckner 1997; Klein et al. 1999a; Cai et al. 2003). Despite its absence, sister chromatids in the 45S rDNA regions are held together and a general cohesin co-factor, SCC3 (Chelysheva et al. 2005), can be detected there, suggesting that a non-meiosis specific cohesin variant provides sister chromatid cohesion for 45S rDNA. These experiments demonstrate that in plant meiocytes the NORs associate with the nucleolus, evade meiosis specific chromatin modifications, axis establishment and SC formation.

### DNA lesions in rDNA are repaired by C-NHEJ during meiosis

Our experiments revealed that the number of DNA lesions, as measured by RAD51 foci, is significantly lower in 45S rDNA compared to the rest of the genome. This suggests that the 45S rDNA is shielded from the meiotic recombination machinery and indeed, mutants with compromised nucleolus integrity (*hda6-6, nuc2-2*) (Earley et al. 2006; Durut et al. 2014) have significantly more RAD51 foci. It is important to emphasize, that these additional foci are not dependent on SPO11. As an independent read-out for DNA lesions and successful repair we quantified the number of 45S rDNA loci at the end of meiosis II. Wild type microspore tetrads should have eight loci with two FISH signals in each microspore, any unrepaired DSB in the rDNA would lead to an increase of rDNA signals. We found that 45S rDNA integrity is only affected in mutants acting in the C-NHEJ DNA repair pathway (*lig4, mre11-4*), while mutants affecting HR repair did not disturb 45S rDNA repair (*com1, rad51*). Consistent with these results, a mutation that abolishes meiotic DSB formation (*spo11-2-3*) did not alleviate rDNA fragmentation in the NHEJ deficient plants. Taken together, this demonstrates that 45S rDNA is protected from SPO11-mediated DSB formation and that SPO11-independent breaks that occur in the rDNA are shielded from HR and repaired via the C-NHEJ pathway.

In somatic cells (human and plants) massive DNA damage in the 45S rDNA (e.g. by generating a DSB in each repeat) activates HR mediated repair. The damaged rDNA repeats then translocate, (only shown for human cells) to the outer border of the nucleolus and form “nucleolar caps” (Sluis et al. 2015; McStay 2016; van Sluis and McStay 2017). Under these conditions, HR is the preferred DNA repair pathway and NHEJ serves as a back-up (Harding et al. 2015). HR leads to deleterious rDNA repeat losses that can only be alleviated in HR-mutant backgrounds in which NHEJ is activated (Sasaki et al. 2010).

Our results indicate that during meiosis in Arabidopsis few, if any, SPO11-mediated DNA breaks are introduced in the rDNA. This has also been observed in yeast (Pan et al. 2011; Vader et al. 2011). It is possible, even likely, that DNA breaks in the rDNA region are the result of collisions between the transcription and replication machineries (Aguilera and Gaillard 2014). Interestingly, *hda6-6* and *nuc2-2* mutants have a strong increase of RAD51 foci at the 45S rDNA and also a strong increase of rDNA array fusions between chromosomes 2 and 4.

Our findings reveal a novel role for NHEJ during meiosis for repairing breaks in the 45S rDNA while it is shielded from the canonical HR machinery.

### The 45S rDNA is sequestered in an HR-refractory chromatin environment

Our results suggest that 45S rDNA constitutes a local, HR-refractory chromatin environment. To test this prediction, we examined the phenotype of ectopically integrated 45S rDNA. The approach was motivated by previous findings, describing the integration of ectopic rDNA repeats into the nucleolus in somatic cell (human and plants) (Wanzenböck et al. 1997) (Grob et al. 2014). Our ectopic repeats are actively transcribed, associate predominantly with the nucleolus and promote pairing of the corresponding genomic regions. Importantly, the DNA around the ectopic rDNA sites had fewer RAD51 foci and meiotic recombination was significantly reduced at two of the sites that reside in recombination-competent regions (Choi et al. 2013). A third ectopic rDNA site (ErDNA5) displayed the same characteristics as the other sites but was positioned near a known CO cold spot and did not show a decrease in meiotic recombination frequency (Choi et al. 2013). We conclude, that 45S rDNA constitutes a local HR-refractory chromatin environment that can repress COs in regions that are normally recombination-competent.

Our results also suggest that the sequence of 45S rDNA is not itself CO-suppressive, but rather that it becomes suppressed when it is sequestered in the nucleolus and acquires a unique chromatin signature. Our study thereby provides a comprehensive answer to the question of how the repetitive rDNA arrays are faithfully inherited.

### List of abbreviations

Double strand breaks: (DSBs)
Non-homologous End Joining: (NHEJ)
Homologous recombination: (HR)
Ribosomal DNA: (rDNA)
Ribosomal RNA: (rRNA)
Fluorescence *in situ* hybridization: (FISH)
Fluorescent Tagged Line: (FTL)
Ectopic ribosomal DNA: (ErDNA)
Artificial pond water: (APW)
Synaptonemal complex: (SC)

## Acknowledgements

We thank all members of the ITN Program “COMREC” for discussions and especially Paul Franzs, Chris Franklin and Monica Pradillo for sharing advanced techniques in cytology. We are grateful to Josef Loidl and Marie-Therese Kurzbauer for critically reading and improving the manuscript. We thank Bernd Edlinger for transforming Arabidopsis plants with the “R4” construct, Maja Sljuka and Teresa Wöhrer for help with the FISH experiments. We also thank Mona von Harder and Zsuzsanna Orbán-Németh for cloning and purification and Katja Schneider for testing the guinea pig anti-ASY1 antibody. We are indebted to Julio Saez-Vasquez for sharing seeds of the *nuc2-2* mutant line, Craig Pikaard for sharing seeds of the *hda6-6* and *atxr5-atxr6* mutant lines and Ortrun Mittelsten Scheid for providing BAC stocks and seeds of the C2R plant line. We also thank the Vienna Biocenter Core Facility for generating the CAS9 constructs and transforming the insect cells for the ASY1 protein expression. We thank the European Union (FP7-ITN 606956) and the Austrian Science Fund (SFB F34; I 3685-B25) for funding.

## Materials and Methods

### Arabidopsis growth conditions

*Arabidopsis thaliana* seeds were stratified in water in the dark at 4°C for two days before sowing on soil/perlite 3:1 mixture (ED 63, Premium Perlite). Pots were covered with a transparent lid until cotyledons were fully developed and first primary leaves visible. Plants were grown under long day conditions in controlled environment rooms (16 hours of light, 8 hours of darkness, 60-80% humidity, 21°C, 15550 lux light intensity) or in short day conditions (8 hours of light, 16 hours of darkness, 60-80% humidity, 21°C, 15550 lux light intensity).

### Selection of transgenic lines

*Arabidopsis thaliana* plants were transformed by the floral dip method (Clough and Bent 1998). Transgenic plants carrying a BASTA resistance gene were selected from a population by spraying the respective herbicide (BASTA^®^, 1:750 in dH_2_O, Bayer CropScience AG (BCS), Monheim am Rhein, Germany) when plants had at least two true leaves. Spraying was repeated every other day until survivors were clearly distinguishable from non-resistant plants. Transgenic lines carrying the Venus-YFP gene under the At2S3 seed storage promoter were selected under a fluorescence stereo-microscope by collecting seeds positive for YFP fluorescence.

### Generation and characterization of *Arabidopsis* ErDNA lines

*Arabidopsis* plants were transformed via the floral dip method with *Agrobacterium* cells containing the R4 binary vector. Transformed plants were selected on MS plates (bacto agar 7 g/L, carbenicillin 500 mg/L, glufosinate ammonium 12 mg/L) supplemented with hygromycin B 30 µg/ml. Resistant plants were transferred from the plates to soil pots and placed in growth chambers. T3 lines that segregated 3:1 were selected as single insertion lines. In order to map the location of the ErDNA insertion, a TAIL-PCR (Thermal Asymmetric Interlaced PCR) was performed by using primers TAIL A2, TAIL B2 and TAIL C2 together with a random primer AD7 as shown in (Liu et al.). DNA fragments of the expected size were sequenced by Sanger sequencing and the integration site was subsequently confirmed by PCR.

### Expression analysis of endogenous rDNA variants and ectopically integrated rDNA repeats

Expression analysis of rDNA variants (length polymorphism in the 3’-ETS) (Pontvianne et al. 2010) was performed by RT-PCR utilizing staged PMCs. Inflorescences from primary and secondary shoots were collected and each bud was dissected individually. Anthers were modestly squashed to force the release of columns (meiotic cells in syncytia). These were collected in artificial pond water (APW: 0.5 mM of NaCl, 0.2 mM NaHCO_3_, 0.05 mM of KCl, 0.4 mM of CaCl_2_ (Miller and Gow 1989)). All meiocytes in prophase I (P) were pooled. Anthers that released single entities (dyads and tetrads) were grouped as meiosis I and II (MM). 20 to 30 anthers per category were collected and frozen immediately in liquid nitrogen. To extract RNA the SV Total RNA extraction KIT (Promega^®^) was used following the product specifications. For reverse transcription of the RNA to cDNA the iSCRIPT cDNA synthesis KIT (BioRad^®^) was used following the product specifications. Finally, in order to amplify the different rRNA/rDNA variants, a specific primer pair (3allrRNAVAR and 5allrRNAVAR) was used, as described in (Pontvianne et al. 2010).

The specific expression of <u>E</u>ctopic <u>rDNA</u> (ErDNA) was determined via RT-PCR, using primers (BstEII-R4Tag-Fw and rDNA-in-rv) directed against a short, integrated stretch of sequence (also including a *Bst*EII site) in the 25S rDNA. The *Bst*EII site served as an ErDNA-specific primer landing platform in order to exclusively amplify rRNA transcribed from the ectopically integrated rDNA repeat(s). In all cases, expression of the ACTIN-7 (AT5G09810) gene served as reference (actin_ampl3_dn and actin_ampl3_up).

### Determination of endogenous and ectopic rDNA repeats numbers

DNA was extracted by crushing leaves in UREA buffer (0.3M NaCl, 30 mM TRIS-Cl pH 8, 20 mM EDTA pH 8, 1% (w/v) N-lauroylsarcosine, 7 M urea) and subsequently purified with Phenol:Chloroform:Isoamylalcohol (25:24:1). The qPCR reaction was performed using the KAPA SYBR FAST kit following the product specifications. To quantify 45S rDNA copy numbers, 20-30 ng of genomic DNA were used together with one primer pair (18SRealdn and 18SRealup) as previously described for *A*. *thaliana* (Muchova et al. 2015). The amplification of the rDNA and of the reference gene ACTIN-7 was performed in separate wells and in three technical replicates. The analysis was conducted with four separate biological replicates. The conditions used for the qPCR were 95 °C 1 minute initial denaturation, 95 °C 30 sec., 55 °C 30 sec., 72 °C 30 sec. for 40 cycles with fluorescence detection after every elongation step. The PCR products were not longer than 250 bp and contained a GC content of approximately 50%. The experiment was performed on an Eppendorf Realplex 2 Mastercycler. Rosette leaves of different sizes, representing different ages, were collected from bottom to top as follows: t0 = 3,5 cm leaves, t1 = 2,5 cm, t2 = 1,5 cm, t3 = 1 cm for short day conditions.

For the plant lines containing the ectopic rDNA repeats, those with single insertion sites in the genome were selected according to their segregation patterns. The copy numbers of integrated rDNA repeats at these single insertion sites were determined by qPCR (again taking advantage of the short, integrated stretch of sequence including a *Bst*EII site in the 25S rDNA). In order to normalize the qPCR results and have an absolute quantification of gene copy number we used primers (HPT_up_reverse and HYG_down) hybridizing to the hygromycin resistance gene and the previously characterized C2R line (Mittelsten Scheid et al. 2003) that harbours only a single T-DNA at a single integration site.

### Preparation of pollen mother cells (PMC) DAPI spreads

Inflorescences were harvested into fresh fixative (3:1 96% ethanol (Merck) and glacial acetic acid) and kept over-night (O/N) for fixation. Once the fixative has decolorized the inflorescences, they were placed in fresh fixative (can be stored for over a month at −20 °C) and subsequently one inflorescence was transferred to a watch glass. The yellow buds were removed to collect only white and transparent buds. The white buds were separated from the inflorescence and grouped according to size. This step is necessary in order to obtain preparations with separated meiotic stages.

Afterwards, the buds were washed 3 times with citrate buffer (0,455 ml of 0.1M citric acid. 0,555 ml of 0.1M tri-sodium citrate in 10 ml of dH2O) and submerged in an enzyme mix (0.3% w/v cellulase, 0.3% w/v pectolyase in citrate buffer). Each bud has to be submerged in order for the digestion to work efficiently. The buds were incubated for 90 min in a moisture chamber at 37 °C. Digestion was inhibited by adding cold citrate buffer (buds can be kept O/N at 4 °C). At this point the buds were transferred (max 3-4 buds of the same size) to a glass slide. Excess liquid was removed and 15 µl of 60% acetic acid added. The buds were suspended by using a metal rod and an additional 10 µl of 60% acetic acid was added to the suspension. The droplet area was labelled using a diamond needle and fixed with fixative 3:1. Slides were dried for at least 2 hours. In order to stage the meiocytes, 15 µl of 2 µg/ml 4’,6 diamidino-2-phenylindol (DAPI) diluted in Vectashield (Vectorlabs) was added to the slide and sealed with a glass coverslip. Images were taken on a Zeiss Axioplan microscope (Carl Zeiss) equipped with a mono cool-view CCD camera. (Vignard et al. 2007)

### Fluorescence *in situ* hybridization and detection of DNA-RNA hybrids

The DAPI slides selected for fluorescence *in situ* hybridization (FISH) were washed in 100% ethanol until the coverslips could be easily removed (5-10 min) and subsequently washed in 4T (4X SCC and 0.05% v/v Tween20) for at least 1 h in order to remove the mounting medium.

After washing the slides in 2X SCC for 10 min they were placed in pre-warmed 0.01 M HCl with 250 µl of 10 mg/ml Pepsine for 90 seconds at 37 °C. The slides were then washed in 2X SCC for 10 min at room temperature. 15 µl of 4% paraformaldehyde (PFA) were added onto the slides, covered with a strip of autoclave bag and placed for 10 min in the dark at RT. The slides were then washed with deionized water for 1 minute and dehydrated by passing through an alcohol series of 70, 90, 100 %, for 2 minutes each. Slides were left to air-dry for 30 min.

Meanwhile, the probe mix was prepared by diluting 1 µl of probe (2-3 µg of DNA) in a total of 20 µl of hybridization mix (10% dextrane sulphate MW 50,000, 50% formamide in 2x SSC). In case the rDNA LNA probe was applied, only 50 pmols (final concentration) were used per slide. The probe mix was denatured at 95 °C for 10 min and then placed on ice for 5 min. Afterwards, the probe mix was added to the slide, covered with a glass coverslip, sealed and placed on a hot plate for 4 min in the dark at 75 °C. Finally, the slides were placed in a humidity chamber over-night at 37 °C. After hybridization, the coverslips were carefully removed and the slides were treated with 50% formamide in 2X SCC for 5 min in the dark at 42 °C. The slides were then washed twice with 2X SCC for 5 min in the dark at room temperature.

In order to detect DNA:RNA hybrids, the Kerafast antibody [S9.6] (1:50 dilution in blocking solution: 1 % BSA, 0.01 % NaN_3_ (w/v) in 1x PBS) was added at this step to the slides and incubated for 30 minutes at 37 °C. The slides were then washed once for 5 min in 2x SCC and an anti-mouse Alexa 555 antibody (1:400 dilution) was added for 30 min at 37 °C. At this point the slides were washed one additional time in 2X SCC.

Finally, 15 µl of DAPI-Vectashield solution were added to the slide and sealed with a coverslip. Images were taken on a Zeiss Axioplan microscope (Carl Zeiss) equipped with a mono cool-view CCD camera.

### Immuno-FISH (<u>T</u>argeted <u>A</u>nalysis of <u>C</u>hromatin <u>E</u>vents – TACE)

Inflorescences with at least two open flowers from the primary shoots of the plant were collected and placed in a petri dish with wet filter paper.

The buds were dissected with needles under a dissection microscope and all anthers with transparent lobes were transferred to a droplet of artificial pond water (APW; 0.5 mM NaCl, 0.2 mM NaHCO_3_, 0.05 mM KCl, 0.4 mM CaCl_2_ in deionized water) on a charged glass slide. Afterwards the anthers were squashed between the tips of dissection forceps to release all meiocytes grouped in syncytia (columns).

The slide was transferred to a light microscope and columns were collected into a 1.5 ml tube on ice with a glass capillary(Chen et al. 2010). 15µl of digestion solution (1 % cytohelicase, 1.5 % sucrose and 1 % polylvinylpyrollidone) was added to the meiocyte solution and carefully mixed. The tube was incubated for 10 min in the dark at room temperature.

Digestion was stopped by putting the tube on ice and 8µl of digested meiocytes were transferred to a clean and charged microscope slide. 20 µl of 2 % v/v Lipsol was added to the droplet and mixed by tilting the slide. After 4 min at room temperature, 24 µl of 4% formaldehyde were added to the slide and left to air-dry completely. (Kurzbauer et al. 2012) Dried slides were washed in 2x SSC for 5 min and the primary antibody mix was added after removing excess liquid. The slides were covered with a piece of autoclave bag and incubated in a humidity chamber at 4 °C over night.

The autoclave bag cover was removed, the slides were washed in 2X SSC for 5 min and the appropriate secondary antibody mix was added. The slides were again covered with an autoclave bag and incubated in a humidity chamber for 1 h at 37 °C.

The slides were washed for 5 min in 2X SCC and 15 µl of 4% formaldehyde fixing solution was added to the slide, covered with a strip of autoclave bag and incubated for 10 min in the dark.

Afterwards, slides were rinsed by dipping into deionized water and dehydrated by passing through an alcohol series of 70, 90, 100 % EtOH, for 2 min each. Finally, the slides were left to dry completely for 30 min in the dark.

In order to prepare the probe mix, 14 µl of hybridization solution was added to 2-3 µl of BAC probes (2-3 µg) or 1 µl of 1 µmol LNA probe and filled up to a final volume of 20 µl with deionized water.

The probe mix was denatured for 10 min at 95 °C and then placed on ice for 5 min. After cooling, the probe mix was added to the slide and covered with a glass coverslip. The borders of the cover slip were sealed with rubber cement and placed on a hot plate for 4 min in the dark at 75-80 °C (75 °C for repetitive DNA and 80 °C for multiple BACs).

The slides were incubated in a humidity chamber O/N at 37 °C. If an LNA probe was used, 3-4 hours were sufficient for a successful hybridization.

Thereafter the slides were washed in 50% formamide-2X SCC for 5 min in the dark at 42 °C and twice in 2X SCC for 5 min.

Finally, 15 µl of DAPI-Vectashield were added to the slide and covered with a glass cover slip. The borders of the glass slide were sealed with transparent nail polish. Slides were imaged with a conventional fluorescence microscope (Zeiss Axioplan). Z-stacks with 100 nm intervals were acquired, deconvolved using AutoQuant software and are presented as projections done with the HeliconFocus software. Super-resolution images were acquired using the Abberior STEDYCON system.

All BACs were labelled by using the Nick Translation mix from Roche following the manufacturer’s instructions. Each BAC was labelled individually and concentrated in a tube. In this study, fluorescently labelled nucleotides Chromatide Alexa Fluor 488-5-dUTP (Thermo Fisher), Chromatide Texas Red-12-dUTP (Thermo Fisher) and Cy5-dUTP (GE Healthcare) were used.

The dilutions used for the different primary and secondary antibodies are as follows: anti-H3K27me1 1:200, anti-H4(Ac) 1:50, anti-ASY1 1:10000, anti-RAD51 1:300, anti-ZYP1 1:500, anti-SCC3 1:500, anti-REC8 1:250, anti-guinea pig Alexa488 1:400, anti-rabbit Alexa 555 1:400, anti-rat 555 1:300, anti-mouse Alexa 555 1:400, anti-guinea pig STAR 580 (for STEDYCON only) 1:500, anti-rabbit STAR RED (for STEDYCON only) 1:250.

### <u>Who</u>le <u>M</u>ount Immuno-<u>FISH</u> (Who-M-I FISH)

This method is an adapted version of the “Whole Mount FISH in a Tube” from Bey et al 2018. Inflorescences were collected from the primary shoots of an Arabidopsis plant and all open flowers and pollen-containing buds were removed with forceps and a needle under a dissection microscope.

All remaining buds were opened with the help of two needles in order to increase accessibility of anthers to the enzyme solution and the FISH probe. The inflorescences were placed in 500 µl fixation solution (1% formaldehyde, 10% dimethylsulfoxide, 1X PBS, 60 mM EGTA) and all buds were submerged.

The solution with the buds was placed under vacuum for 10 min at room temperature and incubated for further 30 min at room temperature.

After removing the fixation solution the samples were incubated 2 × 10 min in 500 µl methanol.

Methanol was removed and samples were incubated for 2 × 5 min in 500 µl ethanol.

Finally, after the ethanol was removed the samples were incubated in 500 µl xylene/ethanol (1:1) for 15 min. The samples were transferred to a new tube with 500 µl of xylene and incubated for 30 min at 50 °C. Samples were washed twice in 500 µl of ethanol and 500 µl of methanol and washed three times in 500 µl of PBT (1X PBS with 0.01% v/v Tween 20).

The samples were digested by incubating the buds in the enzyme solution (0.6% cytohelicase, 0.6% pectolyase, 0.6% cellulase in citrate buffer, pH 4.5) for 1 hr at 37 °C.

The samples were gently washed twice in 500 µl 2X SSC and then incubated in 500 µl of 0.1 mg/ml of RNaseA in 2X SSC for 1 hr at 37 °C. Afterwards the samples were washed twice in 500 µl 1X PBT and fixed in 1% formaldehyde (in 1X PBT) for 30 min.

Samples were then washed twice in 500 µl 1X PBT and once in 500 µl 2X SSC, followed by incubation in 500 µl of a 1:1 mix of HB50 (50% formamide, 2X SSC, 50mM NaH_2_PO_4_) and 2X SSC for 30 min, incubated in 500 µl HB50 for 30 min and finally incubated in 30 µl of hybridization solution (labelled DNA probe 0.5-1.5 µg for DNA repeats and 2-5 µg for unique sequences, 50% formamide in 2X SSC) in the dark for 1 hr.

In order to denature the probe and the target sequence the tube was placed in a heating block for 4 min at 85 °C and afterwards on ice for 3 min. Finally, to hybridize the labelled probe to the sample the tube was left overnight in the dark at 37 °C.

The samples were washed in 500 µl HB50, incubated in fresh 500 µl HB50 at 42 °C for 1 hr and washed in 500 µl PBT for 2 times 10 min each.

In order to successfully immune-label proteins in a whole mount preparation, once the samples were washed in PBT, they were incubated in 50 µl of primary antibody mix under vacuum for 30 min and then transferred for 3 hours to 37°C or overnight at 16°C. The primary antibody mix was 10 times more concentrated than what is usually used for regular spreads. Afterwards, the samples were washed four times in 1X PBT and incubated with 50 µl of the secondary antibody solution for 30 min under vacuum and placed for 3 hr at 37 °C. Finally, the sample was washed three times for 15 min in 1X PBT, incubated in 100 μl 1X PBS with 8μl of DAPI 5 μg/ml for 30 min and placed on a glass slide with 20 µl DAPI-Vectashield. Imaging of whole-mount samples was performed with a Zeiss LSM710 equipped with an AiryScan Unit with 160 nm resolution in x, y and z. The images were deconvolved and 3D rendered with the Huygens Software.

### Scoring recombination rates

Recombination analysis was performed according to the protocol published by (Berchowitz and Copenhaver 2008). In brief, individual flowers were collected from the main shoot and tapped on a glass slide with pollen sorting buffer (10 mM CaCl_2_, 1 mM KCl, 2 mM MES and 5% (w/v) sucrose, pH 6.5) (Yelina et al. 2013) for 2-3 minutes until all pollen was released. Tetrads were scored with an inverted microscope equipped with three filter sets for GPF, RFP and CFP. All plant lines used for tetrad analysis were in the *qrt1-2* mutant background.

### CRISPR-Cas9-mediated ErDNA deletion

In order to specifically delete the rDNA portion of the R4 transgene (Wanzenbock et al. 1997) but retain the resistance marker and the Left Border (LB) and the Right Border (RB) of the vector after integration in the plant genome, a CRISPR Cas9 vector was constructed as described before (Richter et al. 2018). Two gRNAs were designed that bind between the LB of the R4 binary vector and the start of the rDNA sequence and between the 3’ETS and the promoter of the hygromycin resistant gene, respectively. All gRNAs were designed to contain at least 50% GCs and to be next to a PAM sequence. The plants transformed with the Cas9 construct were selected with BASTA and the survivors evaluated for Cas9 activity on both target sites. This was achieved by amplifying by PCR the gRNA target regions within the T-DNA, followed by subsequent Sanger sequencing.

Only T1 plants that showed Cas9 activities on both target sites were propagated to the T2 generation and genotyped for loss of the ectopic rDNA unit within the transgene. It was necessary to go through at least two plant generations to select for a homozygous deletion and for loss of the Cas9 transgene.

## QUANTIFICATION AND STATISTICAL ANALYSIS

### Definition and quantification of RAD51 foci

Related to Figures 4, S4 and 6. RAD51 foci were counted manually on deconvolved, slice aligned and 16-bit projected images. Only RAD51 foci colocalizing with the chromatin (DAPI) were accepted. Only RAD51 foci overlapping with 50% or more with the genomic region specifically labelled with a FISH probe were scored as a colocalizing event.

### Quantification of rDNA fragmentation via FISH

Related to Figure 5 and S5. rDNA fragments were counted manually on 16-bit images of meiotic cells from metaphase II to tetrad stage. Only rDNA signals (FISH) that overlapped with a chromatin signal (DAPI) were accepted.

### Nucleolus-association of the ErDNA

Related to Figure 6 and Video 6. Who-M-I-FISH images were 3D reconstructed in order to locate the nucleolus as an empty pocket within the nucleus. The specific genomic regions (FISH signals) were accepted as “nucleolus-associated” in case they were located at the borders or within the nucleolus itself. All experiments were performed by also detecting the endogenous 45S rDNA as a reference for the nucleolus location.

### Statistical Analysis

t-tests, Mann Whitney test and binary logistic regression tests (as indicated in the figure legends) were performed using GraphPad Prism software and http://statpages.org/logistic.html.

Unpaired, two-tailed Mann-Whitney tests were performed, since D’Agostino Pearson omnibus K2 normality testing revealed that most data were not sampled from a Gaussian population and nonparametric tests were therefore required. Error bars indicate standard deviations.

## Figure Legends

**Figure S1:**
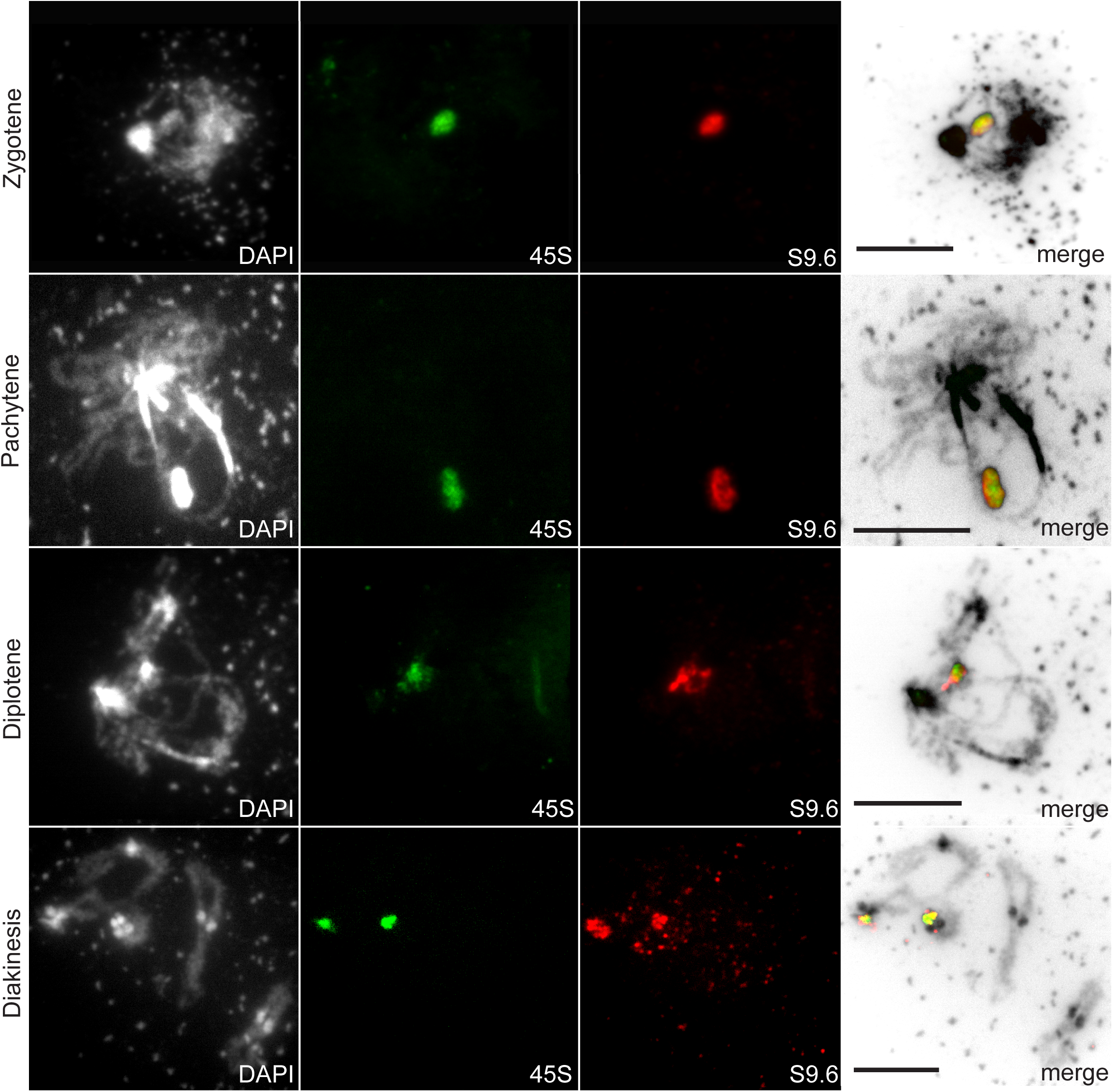
The NORs are highly dynamic regions that are both transcribed during meiosis. Spread nuclei of PMCs from different meiotic stages (as indicated). The S9.6 antibody is directed against DNA:RNA hybrids (red), 45S rDNA has been visualized with a specific FISH probe (green) and DNA has been stained with DAPI (white) (zygotene n=11, pachytene n=12, diplotene n=7, diakinesis n=5). Size bars = 10 µm

**Figure S2:**
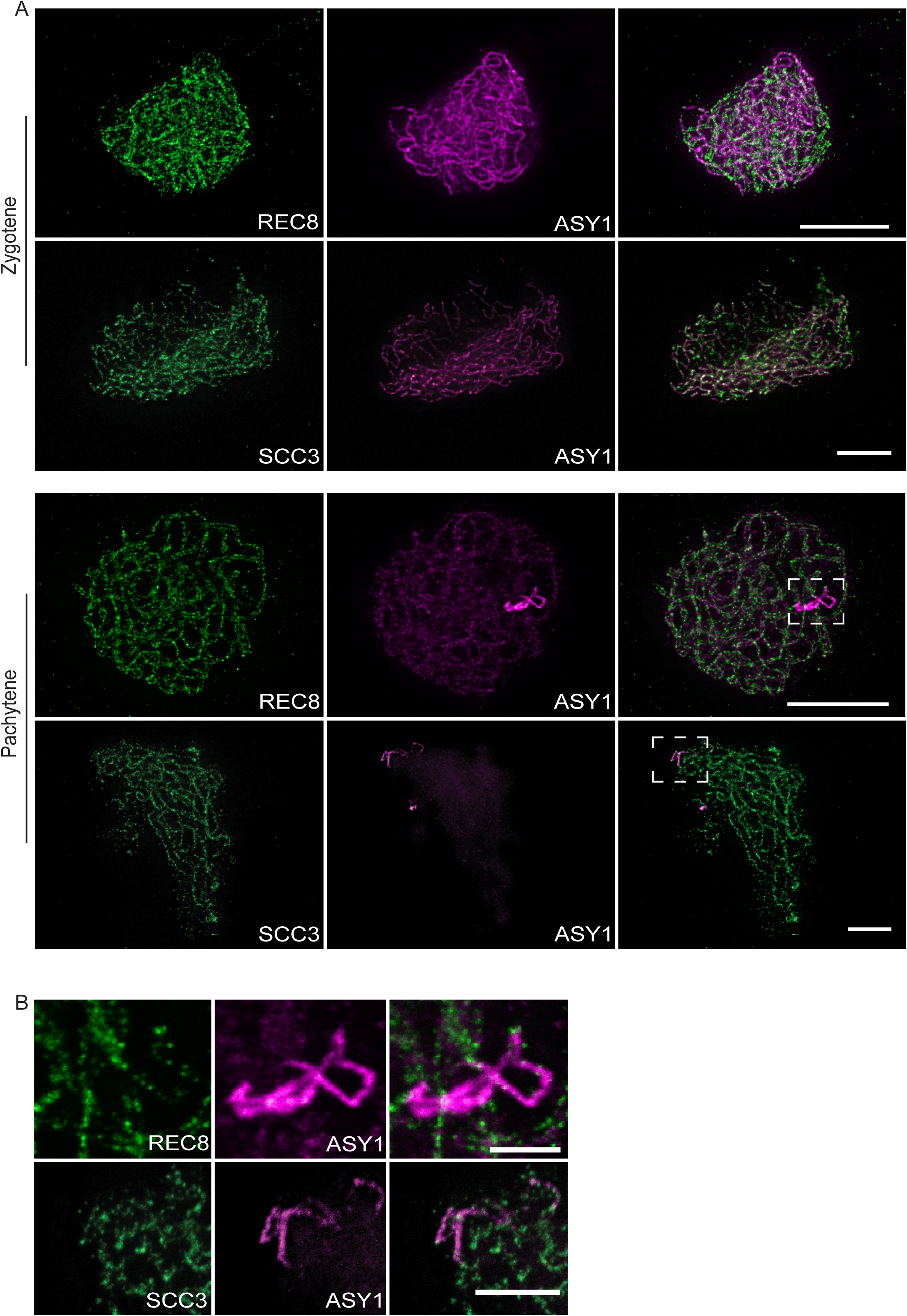
The rDNA acquires distinct chromatin characteristics during meiosis. Super-resolution images of spread meiotic nuclei at zygotene and pachytene stages stained for the axis with an anti-ASY1 antibody (magenta) and for cohesin subunits with anti-REC8 or anti-SCC3 antibodies (green). The nuclei were imaged with an Abberior STEDYCON microscope at 40-60 nm resolution. (A) Individual channels and merged pictures related to Figure 2 are displayed. Size bar = 10 µm. (B) The boxed areas from panel (A) have been enlarged to highlight the localization of cohesin subunits relative to the axis protein ASY1. Size bars = 2 µm.

**Figure S3:**
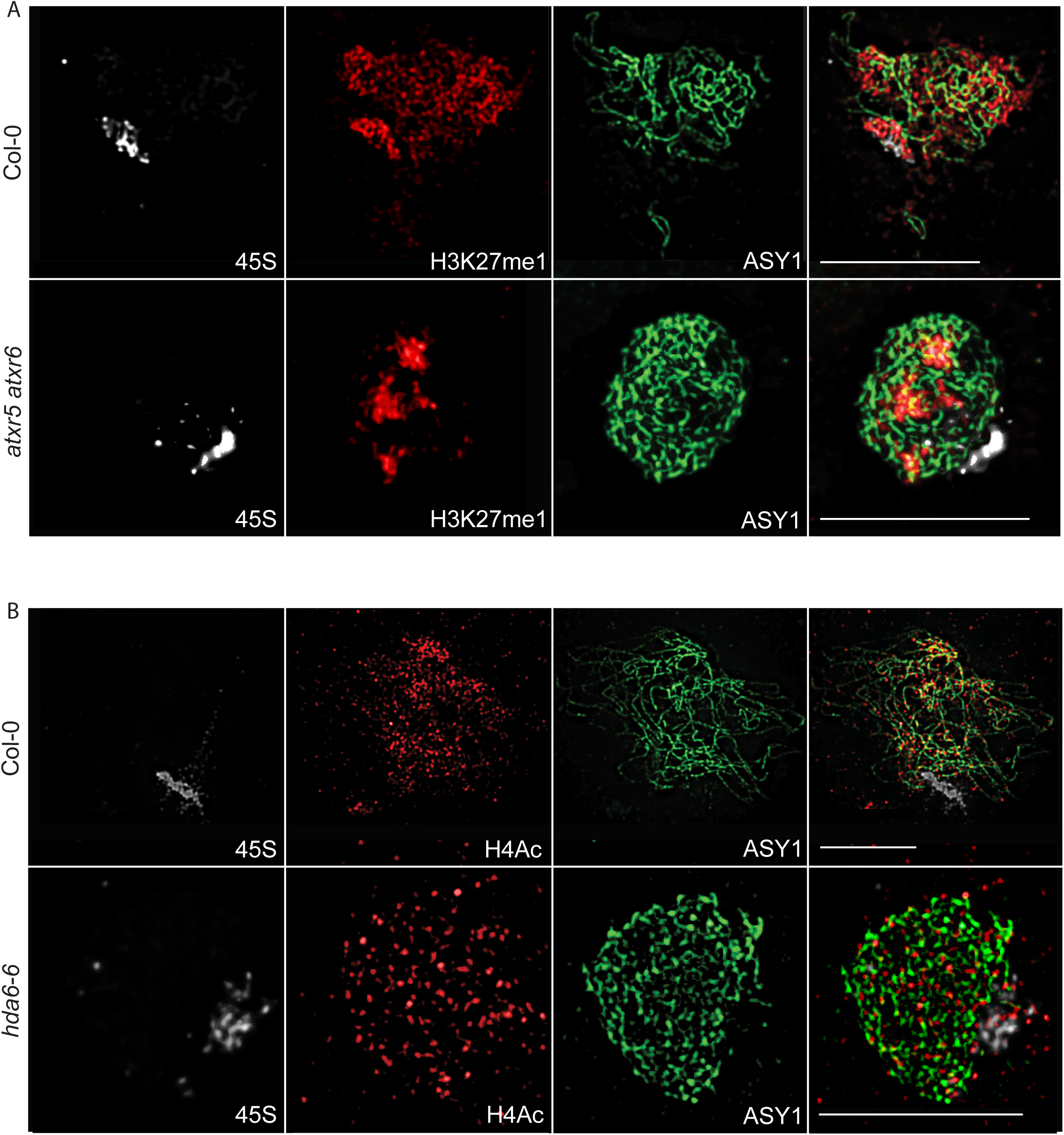
The rDNA acquires a specific chromatin environment during meiosis. (A) Immuno-FISH spreads of wild type (Col-0) and mutant (*atxr5 atxr6*) PMCs at zygotene. The axis has been stained with an anti-ASY1 antibody (green) and the histone modification H3K27me1 with a specific antibody (red). The 45S rDNA has been visualized with a specific FISH probe (white). The histone modification H3K27me1 localizes differently in *atxr5-1 atxr6-1* double mutants compared to Col-0 and does not colocalize with the 45S rDNA. (B) Immuno-FISH spreads of wild type (Col-0) and mutant (*hda6-6*) PMCs at zygotene. The axis has been stained with an anti-ASY1 antibody (green) and the histone modification H4Ac with a specific antibody (red). The 45S rDNA has been visualized with a specific FISH probe (white). The histone modification H4Ac colocalizes with the 45S rDNA in the *hda6-6* mutant background during zygotene in contrast to Col-0. Size bars = 10 µm.

**Figure S4:**
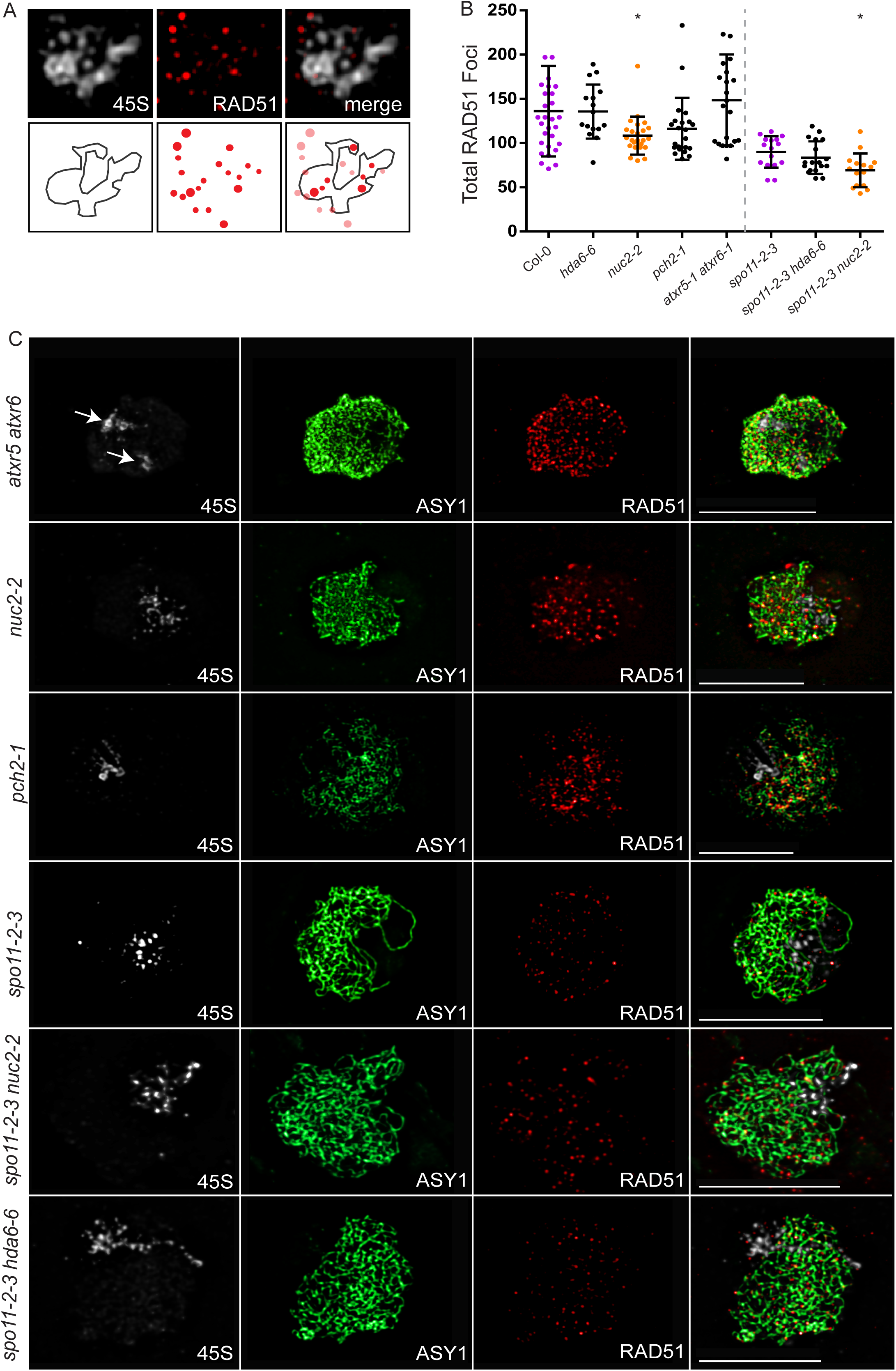
The nucleolus shields rDNA from meiotic DSB formation and deleterious HR. (A) RAD51 – 45S rDNA colocalization analysis. Only RAD51 foci that had an overlap of at least 50% with the 45S FISH signal were counted as colocalizing. (B) Total number of RAD51 foci per nucleus counted at zygotene. Statistical analysis was performed by using a Mann Whitney test. (C) Immuno-FISH spreads of various mutant PMCs (*atxr5-1 atxr6-1, nuc2-2, pch2-1, spo11-2-3, spo11-2-3 nuc2-2* and *spo11-2-3 hda6-6*) at zygotene. The axis has been stained with an anti-ASY1 antibody (green) and the recombinase RAD51 with a specific antibody (red). The 45S rDNA has been visualized with a specific FISH probe (white). White arrows indicate the separated NORs in the *atxr5-1 atxr6-1* double mutant. Size bars = 10 µm.

**Figure S5:**
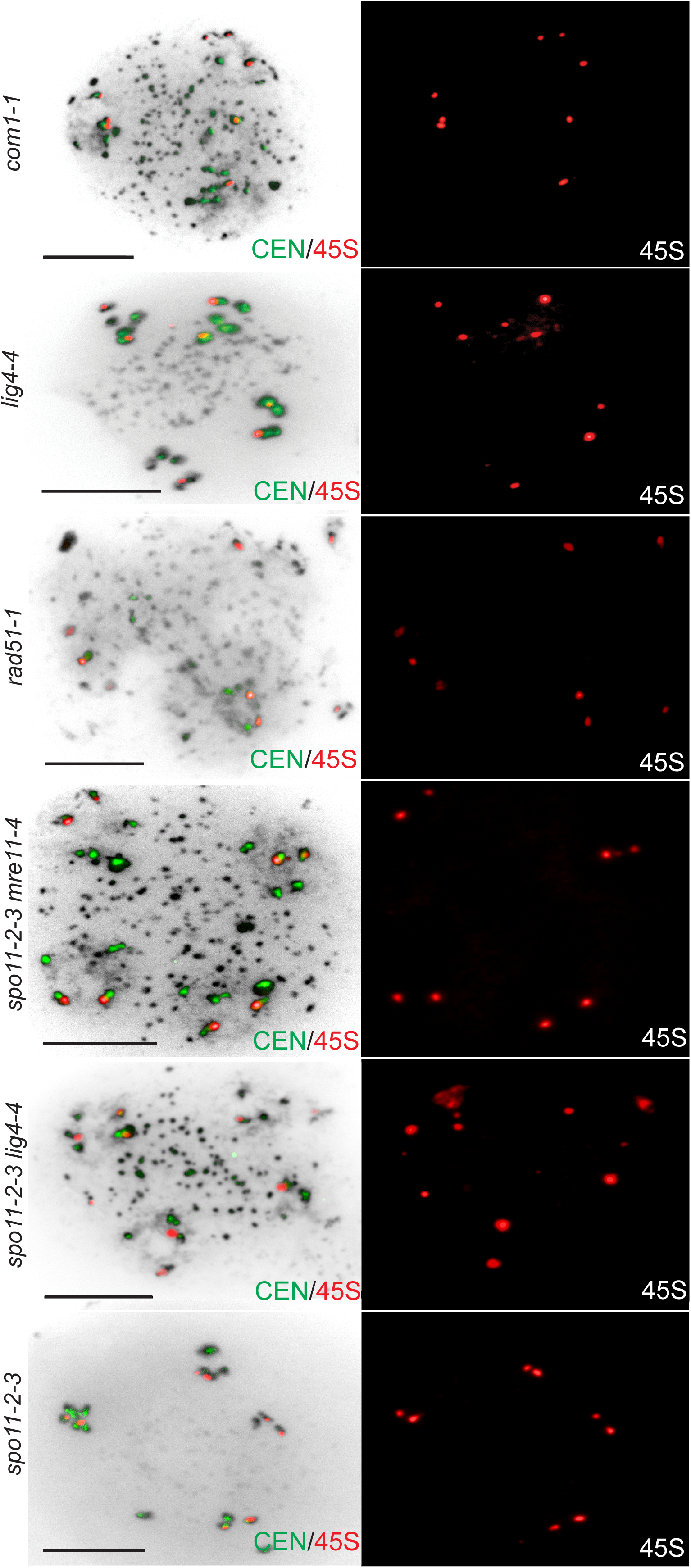
The rDNA is repaired by NHEJ. Spread nuclei of meiocytes from various mutants (*com1-1, lig4-4, rad51-1, spo11-2-3, spo11-2-3 mre11-4* and *spo11-2-3 lig4-4*) at tetrad-like stage followed by FISH to visualize the 45S rDNA (red) and the centromeres (green), combined with panels that only depict the FISH signals that visualize the 45S rDNA (red). More than 8 45S rDNA signals were found in *lig4-4, spo11-2-3 mre11-4* and *spo11-2-3 lig4-4* mutants. Size bars = 10 µm.

**Figure S6:**
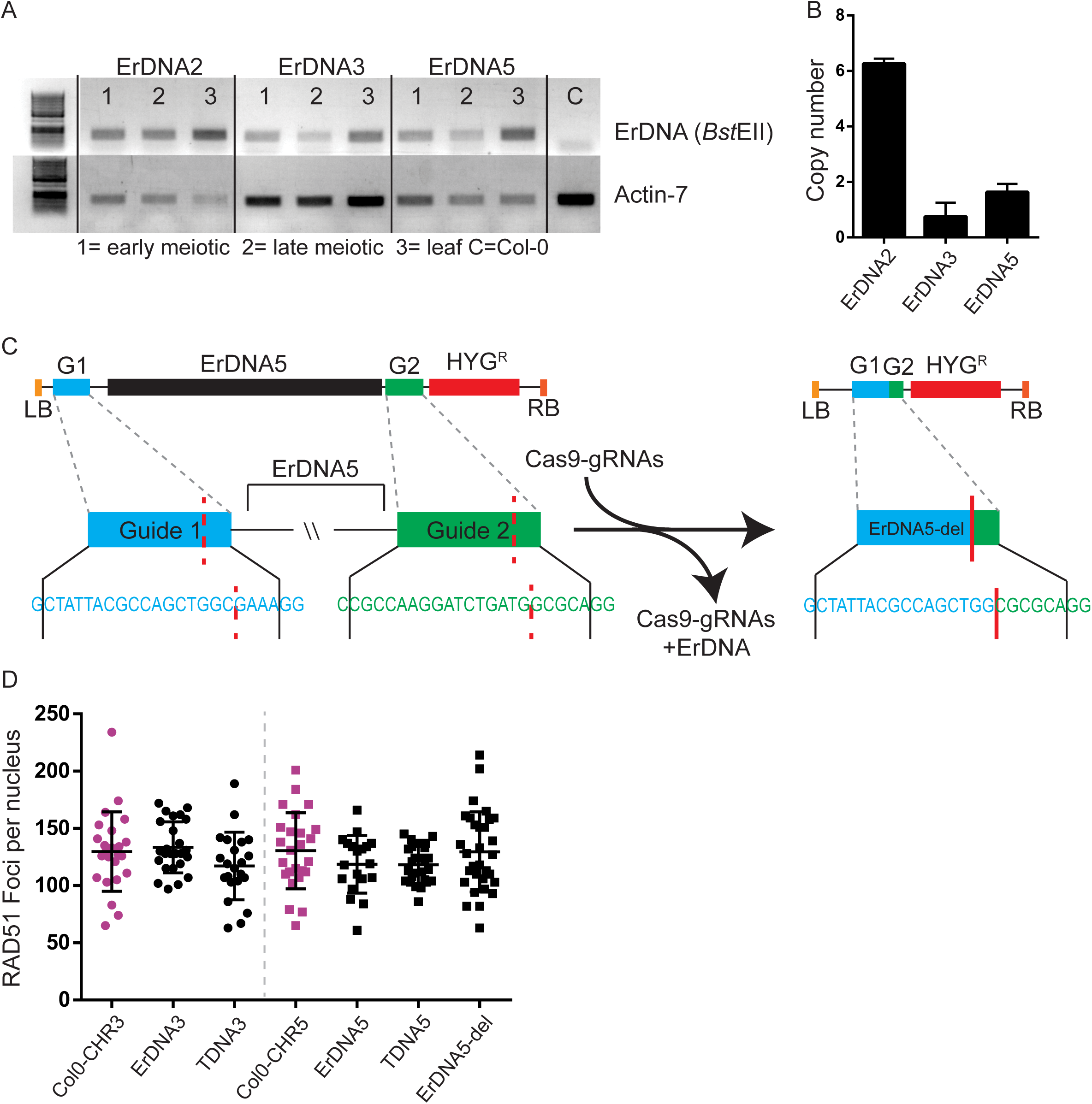
Ectopically integrated rDNA units associate with the nucleolus, promote homologue pairing and suppress meiotic recombination. RT-PCR experiment to test the expression of the ectopic rDNA in plant lines ErDNA2, ErDNA3 and ErDNA5. Total RNA was extracted, reverse transcribed and used as a template for PCR. Utilizing the unique sequence of 20 nucleotides in the ectopic rDNA allowed its specific amplification. Col-0 was used a control plant, not containing an ectopic rDNA unit. The *ACTIN7* gene served as a control against contaminations with genomic DNA. The ectopic rDNA is expressed in all three lines and in all tissues tested (early and late meiocytes, leaves). Ectopic rDNA copy number assessment of the individual ErDNA lines. The *HPT* resistance gene, also encoded by the rDNA T-DNA vector, was used as a further target for quantitative PCR. Normalization was performed utilizing DNA from a plant harbouring a single copy *HPT* transgene for comparison and the endogenous single copy locus *ACTIN7*. (C) Graphical depiction of the ErDNA5 transgene before (left) and after (right) Cas9 activity. GuideRNA/CAS9 target sites are indicated. Sequencing the region of the Cas9 cleavage sites before (left) and after Cas9 activity (right) demonstrated complete deletion of the rDNA portion of the T-DNA insertion. (D) Graph demonstrating that during zygotene in all analysed plant lines total RAD51 foci numbers do not significantly differ from each other (p≥0.8) Statistical analysis was performed by using a Mann Whitney test.

